# SingleM and Sandpiper: Robust microbial taxonomic profiles from metagenomic data

**DOI:** 10.1101/2024.01.30.578060

**Authors:** Ben J. Woodcroft, Samuel T. N. Aroney, Rossen Zhao, Mitchell Cunningham, Joshua A. M. Mitchell, Linda Blackall, Gene W. Tyson

## Abstract

Determining the taxonomy and relative abundance of microorganisms in metagenomic data is a foundational problem in microbial ecology. To address the limitations of existing approaches, we developed ‘SingleM’, which estimates community composition using conserved regions within universal marker genes. SingleM accurately profiles complex communities of known microbial species, and is the only tool that detects species without genomic representation, even those representing novel phyla. Given SingleM’s computational efficiency, we applied it to 248,559 publicly available metagenomes and show that the vast majority of samples from marine, freshwater, sediment and soil environments are dominated by novel species lacking genomic representation (median relative abundance 75.0%). SingleM also provides a way to identify metagenomes for the recovery of novel metagenome-assembled genomes from lineages of interest, and can incorporate user-recovered genomes into its reference database to improve profiling resolution. Quantifying the full diversity of Bacteria and Archaea in metagenomic data shows that microbial genome databases are far from saturated.

## Introduction

A centrally important question asked about microbial communities is determining which microorganisms are present, and at what abundance. The most accurate method for answering these questions involves shotgun metagenomic sequencing of the sample, which generates reads in proportion to the relative abundance and genome size of each community member. These reads are analysed with metagenomic taxonomic profiling software to estimate the relative abundance of each microbial species in the sample.

Metagenomic taxonomic profiling (herein ‘taxonomic profiling’) is typically undertaken by matching reads to databases derived from reference genomes, usually to sets of clade-specific marker genes(Milanese et al. 2019; Blanco-Míguez et al. 2023), kmer matching(Lu et al. 2017; Wood et al. 2019; Irber et al. 2022; Park et al. 2023) or by read mapping to whole genomes(Sun et al. 2023). The most recent version of MetaPhlAn (v4) incorporated a large set of metagenome-assembled genomes (MAGs) into its reference genome database, increasing the fraction of reads it assigned appreciably(Blanco-Míguez et al. 2023). However, this expanded database only includes genomes which are currently assembled and of medium-to-high quality, which means completely new species are missing from the taxonomic profiles MetaPhlAn generates. Taxonomic profiling can also be carried out by matching reads to known protein sequences i.e. a ’BLASTX’. The most widely used tool in this space is Kaiju(Menzel et al. 2016) which classifies reads against all known protein sequences in NCBI nr, Progenomes(Mende et al. 2020), or other large sequence databases.

Despite the wide variety of profiling tools that have been developed and extensively benchmarked, accurate estimation of community composition remains a challenging problem(Meyer et al. 2022; Poussin et al. 2022). Existing taxonomic profiling software is also largely restricted to characterising the abundance of species with reference genomes, missing most novel species. This inability to account for novel species has long been recognized as a central limitation of taxonomic profiling from metagenomic data(Menzel et al. 2016), one that significantly hinders the study of microbial ecology.

Here we present a fast and accurate species-level profiler of short read metagenomes (‘SingleM’) that is able to identify and enumerate lineages where no complete or draft genome exists. It achieves these goals by analysing only those reads which cover highly conserved regions (‘windows’) of single copy marker genes. Restricting analysis in this way structures a metagenomic dataset into a simplified intermediate representation, an operational taxonomic unit (OTU) table for each marker gene. From this representation, new algorithmic approaches can be applied which improve profiling fidelity and open up new possibilities for the interpretation of taxonomic profiles.

## Results and discussion

### Taxonomic profiling through read recruitment to conserved windows

SingleM is a software suite which takes short read metagenomic data as input, and estimates the relative abundance and per-base read coverage of Bacteria and Archaea at each taxonomic level from domain to species (**Figure 1**). SingleM starts by matching reads to highly conserved regions (’windows’) of 59 single copy marker genes (22 Bacteria-specific, 24 Archaea-specific, 13 targeting both domains). Importantly, reads are matched to these conserved gene windows by searching in amino acid space, using DIAMOND BLASTX(Buchfink et al. 2021), maximising recruitment of reads from divergent lineages. This is in contrast to other marker-based taxonomic profilers, which map the nucleotide sequences of reads to markers directly (e.g. MetaPhlAn, mOTUs).

**Figure 1.**
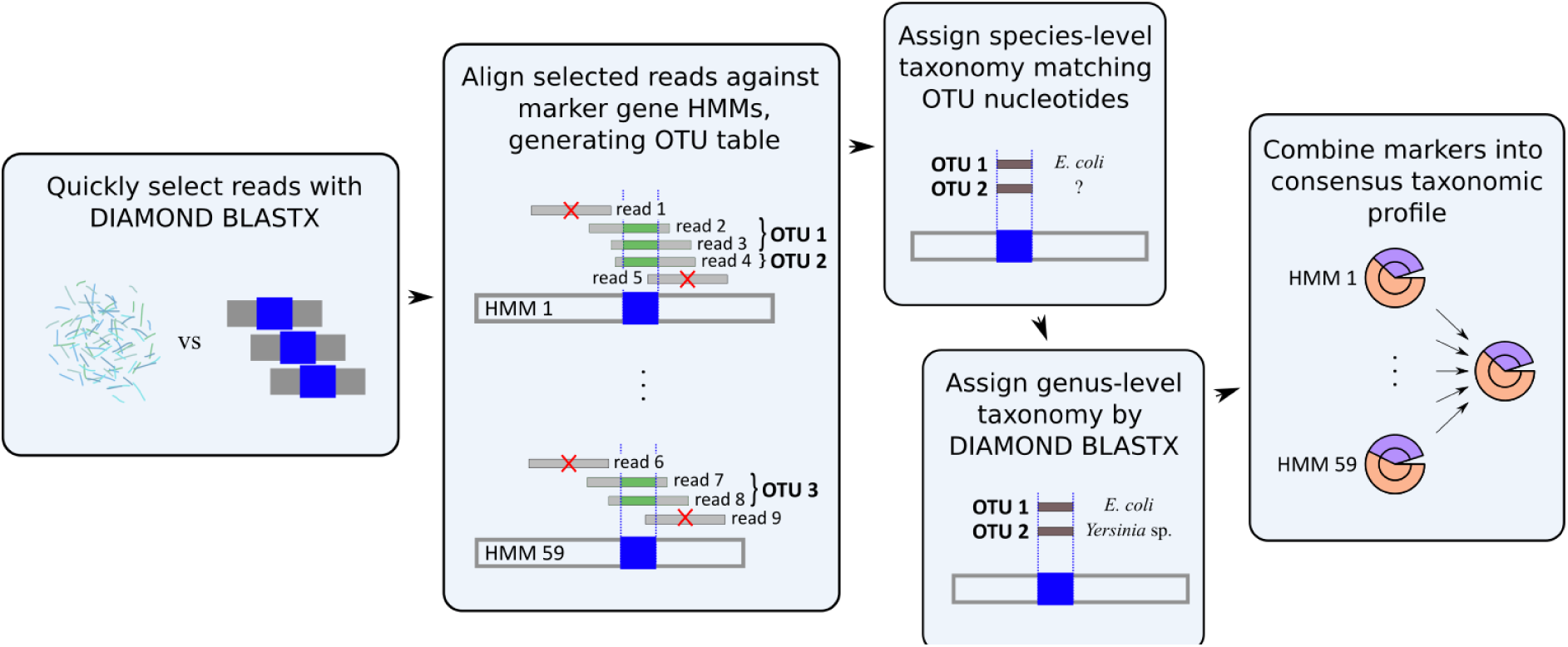
Conceptual overview of the SingleM algorithm. Raw metagenomic reads are first filtered to find those that are homologous to any of the 59 marker genes. Selected reads are translated and aligned to their marker’s hidden Markov model (HMM), discarding any which do not fully cover the 20 amino acid window. The remaining reads are clustered into operational taxonomic units (OTUs) using the corresponding 60 nucleotides. Taxonomy is assigned to each OTU either at the species or genus levels by smafa or DIAMOND BLASTX, respectively. In the final step, the assigned taxonomy of each cluster is used to create a taxonomic profile which summarises the read coverage observed across the 59 marker genes.

In SingleM, only those reads which fully cover these 20 amino acid (60 nucleotide) windows are analysed further. The 60bp nucleotide sequences of each read are clustered *de novo* into operational taxonomic units (OTUs). The result is an intermediate representation of the microbial community, an unannotated OTU table for each marker gene that has been created independent of taxonomy. Its completeness relies only on the BLASTX-based matching approach, which we show below has high fidelity even for novel lineages.

To assign taxonomy to each OTU, SingleM uses the Genome Taxonomy Database (GTDB)(Parks et al. 2022) rather than NCBI taxon strings. This decision was motivated by the taxonomic consistency of the GTDB and its use of the 95% average nucleotide identity threshold to delineate species, which helps establish whether each window sequence represents a new species or one known from the reference database. Taxonomic classification is carried out using a custom alignment algorithm ’smafa’ which aligns each OTU’s 60bp window sequence against 60bp sequences derived from GTDB species representatives(Parks et al. 2022). Compared to general purpose sequence similarity search algorithms, smafa rapidly identifies the most similar sequences without resorting to algorithmic heuristics. This task is made feasible by observing that the query and subject sequences have already been aligned to the marker window and therefore to each other. If no GTDB species encodes the query window sequence within 96.7% average nucleotide identity (**Supplementary Note 1**), then a truncated genus-level taxonomy is assigned using a DIAMOND BLASTX best hit approach.

In the final step, a summarised taxonomic profile of the metagenome is created by integrating the information available for each marker gene. The composition of both known species and higher level taxons is estimated by applying an expectation-maximisation algorithm(Kim et al. 2016) which considers the abundance and taxonomic assignment made to each OTU. Then, to estimate the abundance of each taxon, the abundance of OTUs assigned to the taxon or its descendents are summed, for each marker gene. The abundance of each taxon is calculated as a trimmed mean taken across the marker genes, excluding those with total abundance in the lowest and highest 10% to account for taxonomy misassignment and lineages with reduced genomes that do not encode all marker genes. Noise in the taxonomic profile is also reduced by removing all taxons with a total abundance of less than 0.35X, a threshold developed by application of the algorithm to CAMI 1 benchmarks(Sczyrba et al. 2017) and public datasets (data not shown). In these cases the abundance is re-assigned to a higher level taxon with >=0.35X coverage.

### Comparing SingleM to other taxonomic profilers

The taxonomic profiling accuracy of SingleM was first benchmarked on simulated communities which contained genomes from known species, testing against other tools for which a GTDB R207 reference database was available. Complex microbial communities were modelled after the CAMI 2 ’marine’ benchmark datasets(Meyer et al. 2022). We found the performance of SingleM was superior, at an average of >0.13 better Bray-Curtis dissimilarity than all other tools at the species level (**Figure 2A**). SingleM was also the top-ranked tool in terms of F1 score, false positive rate, Jaccard index, L1 norm error and purity (**Supplementary Data 2**), but similar to other marker-based methods was less performant when genomes were present at lower abundance (**Supplementary Note 2**). We note that for MetaPhlAn and mOTUs, use of an officially supported translation step from NCBI to GTDB taxonomy was required for comparison, which may have adversely affected these tools’ accuracy.

**Figure 2.**
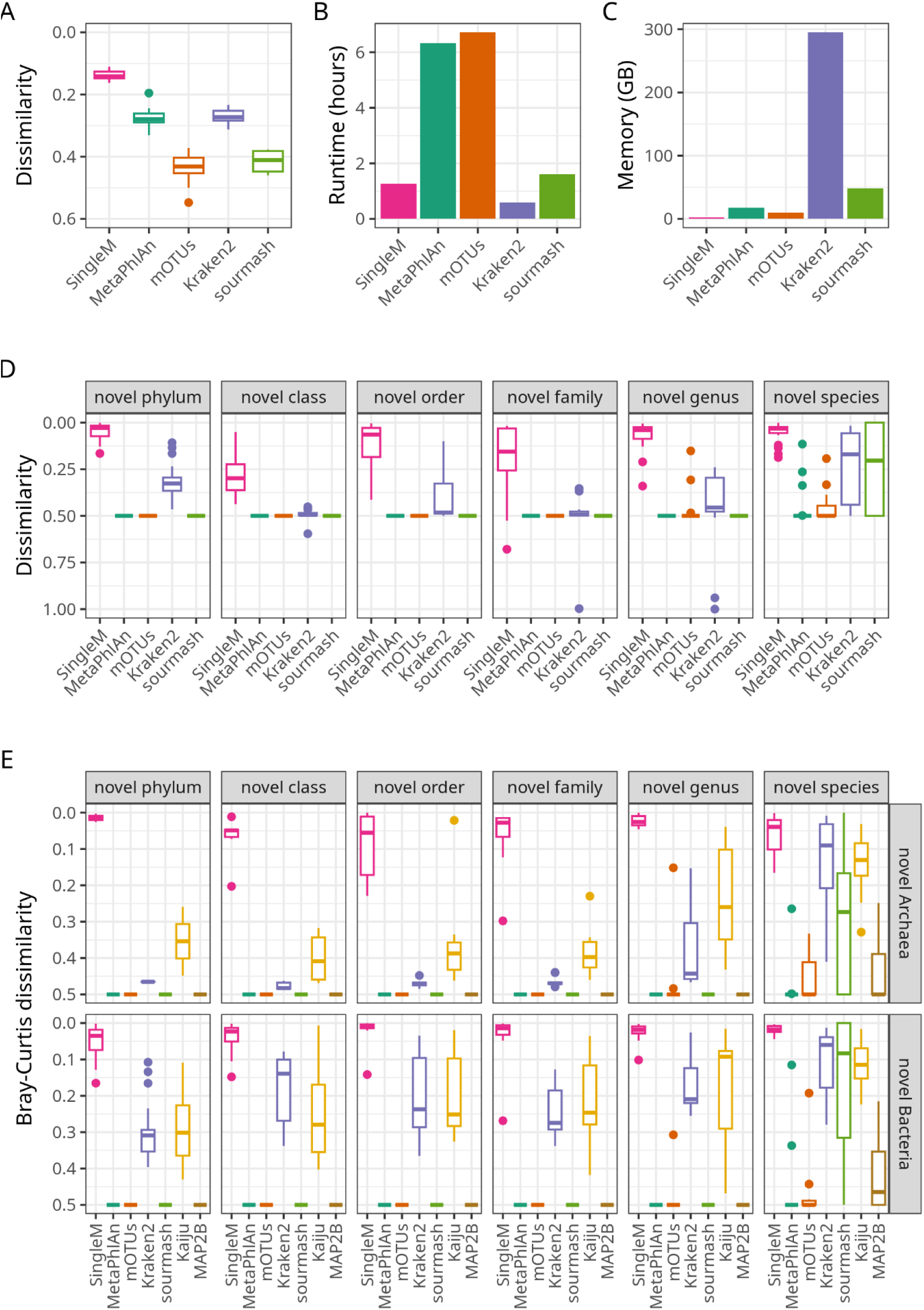
Metagenomic taxonomic profiling tool benchmarks. Complex communities of known species were used to benchmark each tool in terms of (**A**) accuracy defined as Bray-Curtis dissimilarity to true community structure at the species level, (**B**) runtime and (**C**) RAM usage. In (**A**), a dissimilarity of 0 indicates a perfect reconstruction of the mock community. The Kraken2+Backen result is included for context, though this workflow estimates the read count of each species rather than their relative abundance. Kaiju and MAP2B are excluded from the accuracy benchmark as no Genome Taxonomy Database (GTDB) R207 reference database was available. In (**D**), accuracy of each tool is shown for 120 mock communities, where each community was composed of 1 known species and 1 lineage new in R214, at equal abundance. Accuracy was assessed on the most specific rank possible given the constraints of the R207 taxonomy e.g. class level profile dissimilarity for genomes from novel orders. In (**E**), accuracy for each tool is shown for the same communities, but assessed at the kingdom level as a measure of how well each tool detects novel lineages. A Bray-Curtis dissimilarity of 0 indicates full detection, where 0.5 indicates that the novel lineage was completely undetected. Kaiju and MAP2B are included in this benchmark only since they do not output GTDB R207-based taxonomy. In (**D**) and (**E**), the Kraken2+Backen workflow is directly comparable to the other tools since the 1:1 ratio of the two simulated species holds sufficiently for both read count and relative abundance.

In analysing these benchmark datasets, SingleM was fast, using ∼20% of the runtime of MetaPhlAn and mOTUs when using a single CPU, analysing 1.3 million reads per minute (**Figure 2B**). The only faster workflows tested was Kraken2+Bracken, which used 42% of the runtime of SingleM respectively. However, Kraken2+Bracken used a much larger quantity of RAM (295GB). SingleM, in contrast, used the least amount of RAM (2GB). The lightweight runtime requirements of SingleM are a consequence of its optimised upfront detection of reads derived from marker gene windows, such that no further processing of the vast majority of reads is required.

To assess whether SingleM and other profiling tools can accurately represent novel lineages, we selected 120 species which were new in GTDB R214, analysing them with a reference database derived from the previous version R207. For each selected novel genome, reads were simulated at 10X coverage, creating 120 mock communities. To establish a point of reference in these mock communities, a known reference genome from the alternate domain was added at equal abundance i.e. a known bacteria for novel archaea, and a known archaeon for novel bacteria.

The classification accuracy of five profiling tools with available R207 reference databases were assessed by comparing their estimated profiles to the gold standard at the highest resolution possible given the constraints of the R207 taxonomy e.g. class-level Bray-Curtis dissimilarity for genomes from novel orders, order-level dissimilarity for novel families, and so on. On this benchmark, a Bray-Curtis dissimilarity of 0 indicates the gold standard profile was perfectly reconstructed, while 0.5 indicates that the novel lineage was entirely missed by the tool. SingleM showed superior performance across all novelty levels (average 0.13±0.13, **Figure 2D**, **Supplementary Figure 1**) compared to other tools (average 0.46±0.10).

The specific ability of tools to simply detect novel lineages, rather than both detect and classify them, was then assessed using the same benchmark data. Each tool’s ability was assessed by calculating their profile’s Bray-Curtis dissimilarity to the gold standard as before, but at the least resolved taxonomic level possible, the kingdom level. SingleM performed very well in detecting the novel lineages within these 120 mock communities (**Figure 2E**), averaging a Bray-Curtis dissimilarity of 0.04±0.05. In comparison, most other tools scored an average of >0.45 (MetaPhlAn, mOTUs, sourmash, MAP2B). The only exceptions were Kraken2+Bracken and Kaiju, which scored 0.28±0.15 and 0.25±0.14. However, the performance of Kraken2+Bracken and Kaiju on novel archaea was substantially worse (0.38±0.15 and 0.30±0.14) than on novel bacteria (0.21±0.11 and 0.22±0.12). This suggests that their performance on novel bacteria may be partially a consequence of there being more bacterial reference genomes than a true ability to generalise to novel lineages. The bias of all tools other than SingleM against detection of novel lineages was pronounced even when the novel species was contained within a known genus. This was particularly true for previous marker-based methods. We attribute SingleM’s strong performance on this benchmark to its use of a sequence similarity search method based on amino acids rather than nucleotides during read recruitment, which allows divergent marker gene sequences to be detected.

We conclude that most taxonomic profiling tools fail to adequately weight novel lineages in their taxonomic profiles, even when the novelty is only at the species level. In contrast, based on these analyses and others carried out on highly reduced symbiont genomes (**Supplementary Note 3**), we found SingleM reliably detects previously unknown lineages even if they are novel at the phylum level.

### Taxonomic profiles of publicly available metagenomes

Having established SingleM as a scalable and accurate taxonomic profiling tool, we applied it to metagenomes at the NCBI SRA(Kodama et al. 2012) that were publicly available in December 2021. Community profiles were derived from 248,559 metagenomes in 17,617 projects comprising 1.3 Pbp of sequencing data, an amount which was ∼3X the quantity annotated by previous rRNA-based efforts(Martiny et al. 2022). Results of this large scale analysis are made available at the ‘Sandpiper’ website (https://sandpiper.qut.edu.au) where taxonomic profiles can be searched based on GTDB R214 taxonomy strings or dataset accession.

This large set of SingleM-derived community profiles allowed us to estimate how much of the worlds’ metagenomes are represented in reference genome databases, and how much is missing (**Supplementary Note 4)**. In light of recent large-scale MAG mining efforts(Almeida et al. 2021; Nayfach et al. 2021; Paoli et al. 2022; Ma et al. 2023; Schmidt et al. 2023), all community profiles were first reassigned taxonomy using a GTDB reference database supplemented with newly mined MAGs. Known species dominated most host-associated datasets, with an average of 78% of each community assigned a species level taxonomy after weighting by relative abundance (**Figure 3, Supplementary Table 1**). A higher average (henceforth ’known species fraction’) was observed in human and mouse metagenomes (80% and 85%), likely due to their being the subject of more studies (111, 297 and 7,354 metagenomes respectively, **Supplementary Data 3**) and comparatively less diverse communities. Bovine, pig and plant-associated metagenomes are less well represented in reference databases (46%, 71% and 56%). In contrast, the known species fraction was much lower in environmental metagenomes. As expected, soils (14%, median 8%) and sediments (20%, median 12%) had the lowest known species fraction. Marine (41%, median 40%) and freshwater (45%, median 46%) metagenomes were somewhat better characterised.

**Figure 3.**
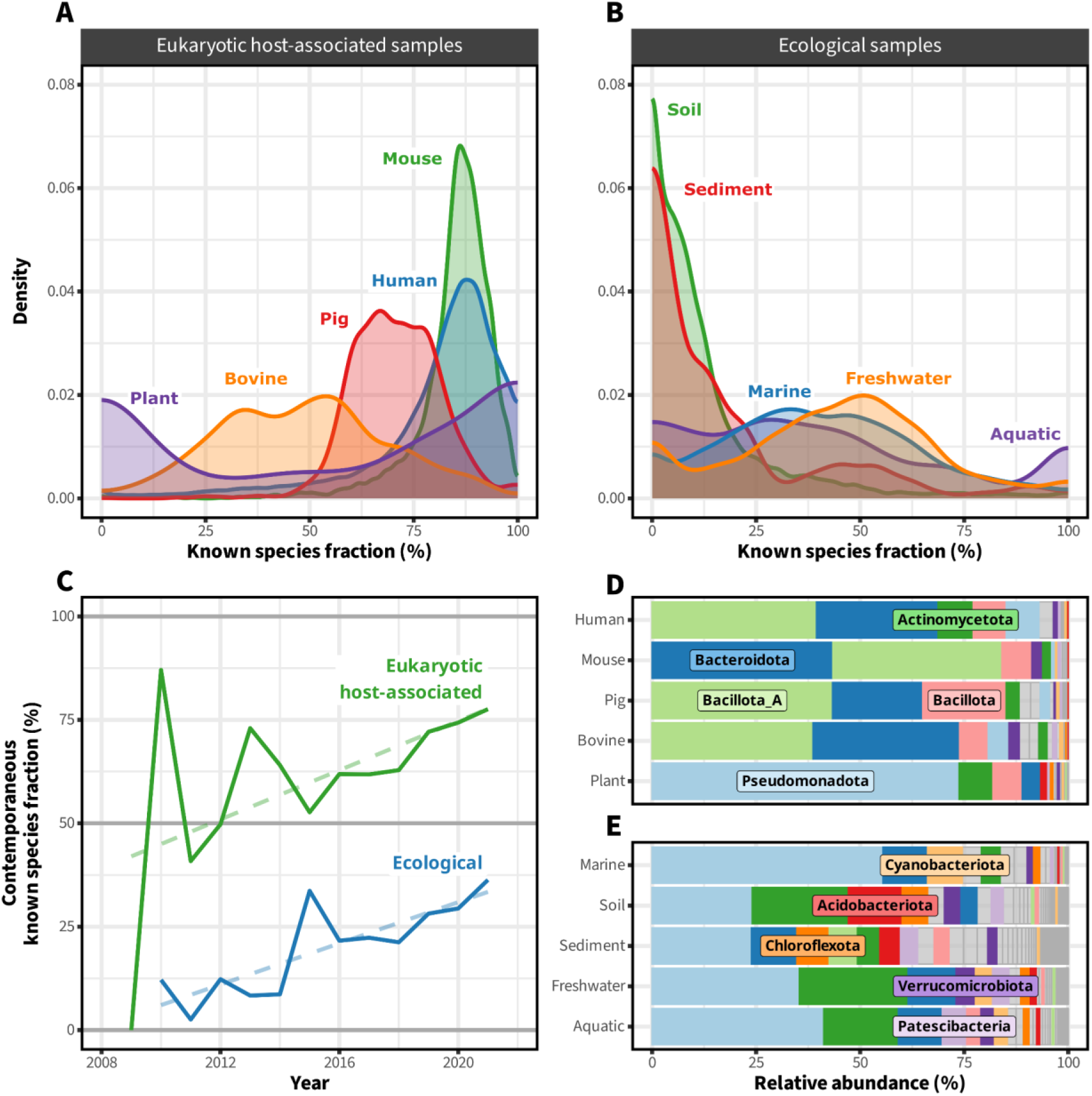
Summary of public metagenomes. Panels (**A**) and (**B**) show the fraction of each metagenome that has been assigned a species-level taxonomy. The remaining fraction currently lacks genomic representation. Plotted is the distribution of these fractions across datasets derived from various eukaryotic host-associated (**A**) and ecological (**B**) environments. In (**C**), the average fraction of metagenomes released in each year assigned to the species level is shown, counting only those species where a genome was available the year before. The dotted line is a linear model of each environment type, weighted by metagenome count. Less metagenomes were published earlier on, so early known species fractions are more variable, spiking as high as 80%. Panels (**D**) and (**E**) show the relative abundance of the phyla observed in selected eukaryotic host-associated and environmental metagenomes, respectively.

Cultured species made up 47% of host-associated taxonomic profiles on average (median 48%, **Supplementary Table 1**). This is consistent with the recent observation that 29% of the UHGG human gut MAG collection has a cultured species representative(Almeida et al. 2021) since higher abundance species are more likely to have been cultured. In contrast, cultured species made up only a very small minority of profiles from marine, freshwater, aquatic, sediment and soil environments (median 2.6%, mean 8.0%). Uncultured species particularly dominated in soils, where a median of 0.8% were cultured (mean 3.5%).

Together, the recent MAG mining efforts added 82,619 new species level lineages to the GTDB R214-based reference database, which was originally composed of 85,205 species. Overall, the median known species fraction in environmental metagenomes was 25.0% (mean 30.2%). However, environmental metagenomes already had a 19.9% median known species fraction prior to the addition of these new MAGs (mean 25.6%, **Supplementary Table 1**). Despite almost doubling the set of available species-level reference genomes, the additional MAGs only improved the median known species fraction of environmental metagenomes by 5.1%. These results underscore the utility of using taxonomic profiling approaches that account for novel lineages and show that a remarkable diversity of organisms are not yet represented in reference genome databases at the species level.

New metagenomic sequencing often detects new microbial diversity, so we next provide a historical view of the rate at which new species are encountered in metagenomic sampling. The average known species fraction of metagenomes released each year was calculated, counting only those species where a genome was available at the start of that year (**Figure 3**). This measure estimates the relative abundance of novel species in newly sequenced metagenomes given the state of the reference database available before sequencing. More than 50% of newly sequenced host-associated metagenomes were assigned at the species level since ∼2012. Steady progress is being made towards high known species fractions in ’ecological metagenomes’ (an NCBI taxonomy category which includes environmental metagenomes and biomes such as wastewater), but at current rates the reference database is much further from saturation.

At the phylum level, Bacteroidota and Bacillota_A (which includes many lineages previously classified as Firmicutes) comprised the majority of commonly sequenced animal metagenomes (human, mouse, pig and cow), with a combined average of 73% (**Figure 3**). Pseudomonadota (previously known as Proteobacteria(Oren and Garrity 2021)) was the most abundant phyla in the 5 most commonly sampled environmental biomes (soil, sediment, marine, freshwater, aquatic), accounting for 36% of average relative abundance. It is also the highest abundance phylum in many less well sampled environments (**Supplementary Data 3**). This phyla also appears frequently in some host-associated metagenomes, dominating plant metagenomes with an average relative abundance of 72%, and ranking in the top five phyla for both pigs and humans. These analyses underline the remarkable ability of Pseudomonadota to adapt to and dominate a wide variety of different environments.

We intend for Sandpiper to be a continually updated resource for the community as new metagenomes are sequenced and genomes recovered. SingleM has largely solved the problem of novel lineage detection (**Figure 2**), so the continual efforts to improve reference databases do not necessitate a full reanalysis of previously processed raw metagenomic reads. Only the taxonomic assignment of OTUs and downstream summarisation into taxonomic profiles need to be recomputed, operations which are markedly less resource-intensive. For instance, updating the 248,559 Sandpiper profiles to GTDB R214 taxonomy only took 2 days and a total of∼30,000 CPU hours on an in-house compute cluster.

### Taxonomically targeted MAG recovery from public metagenomes

One application of the Sandpiper dataset is to inform genome recovery efforts aimed at specific lineages of interest. The assembly and binning of metagenomic datasets involves computationally intensive techniques, making them challenging to apply wholesale to all public datasets. MAG recovery efforts from both human and environmental samples have only been undertaken at the scale of ∼13,000 metagenomes per study(Parks et al. 2017; Almeida et al. 2019; Pasolli et al. 2019; Nayfach et al. 2021; Paoli et al. 2022; Ma et al. 2023), with the exception of the recent SPIRE initiative(Schmidt et al. 2023) (∼100,000 samples). While impressive, these efforts encompass less than half of the metagenomes currently in Sandpiper. Further, improving MAG quality by reapplication of genome recovery pipelines with updated bioinformatic tools requires significant computation. Application of state of the art genome recovery methods across all public datasets is therefore out of most researchers’ reach.

For studies wishing to concentrate analysis on specific taxa, we devised a simple procedure to suggest samples likely to yield novel genomes based upon the estimated coverage and relative abundance of the taxa (see methods). To test the procedure, we attempted recovery of MAGs from four related bacterial phyla, the Muirbacteria, Wallbacteria, Riflebacteria and Fusobacteria. These phyla branch together near the root of Bacteria(Coleman et al. 2021) and are underrepresented in reference databases, with 1, 3, 22 and 95 species representatives available at the time of analysis (GTDB R207), respectively. Further taxonomic sampling of these phyla may inform future efforts to confidently place the Bacterial root.

In this proof of concept experiment, we analysed 63 metagenomes predicted to contain novel species belonging to these phyla at sufficient abundance to enable genome recovery. Novel genomes were successfully recovered from 55 of these metagenomes (87% of samples, 62 MAGs from these phyla in total) with completeness >70% and contamination <5% (average 93% and 2%) (**Supplementary Data 4**). All of these MAGs were novel to at least the species level and include representatives of new genera from each of the four phyla. Genomes from Muirbacteria, Wallbacteria and Riflebacteria phyla were mostly derived from industrial(Yin et al. 2018, 2020; Cheng et al. 2019; Ma et al. 2021) or environmental communities. Recovered Fusobacteria were associated with non-human eukaryotic hosts including insects(Laviad-Shitrit et al. 2020), birds(Cao et al. 2020), monkeys(Rhoades et al. 2021), and fish(Le Doujet et al. 2019; Riiser et al. 2020; Collins et al. 2021; Pratte et al. 2022). We conclude that Sandpiper can be used to expand the diversity of genomes present in reference databases through the targeted application of genome recovery pipelines.

### Supplementing reference data with newly recovered genomes

Genome-centric workflows have become a mainstay of metagenomic analysis due to their ability to recover genomes from samples *de novo*. However, assembly and binning typically only yield MAGs for a subset of community members due to limited coverage or high strain heterogeneity(Meziti et al. 2021). To estimate relative abundance in their microbial communities, researchers are usually forced to restrict analysis to MAGs they themselves recovered, or to use general reference databases that exclude their MAGs. To enable a more holistic taxonomic profile to be obtained in these scenarios, we provide a ’supplement’ mode of SingleM, which adds genomes to the SingleM reference database. Profiling metagenomes with this supplemented reference database enables users to integrate the wealth of data available in reference genome databases with their newly discovered MAGs.

## Conclusion

Single copy marker genes have long been used in microbial ecology for predicting the quality of assembled genomes(Parks et al. 2015), for phylogenomic inference(Wu and Eisen 2008) and for taxonomic profiling(Milanese et al. 2019). Here we have established that not only are entire genes conserved, but specific motifs are sufficiently conserved to allow unassembled reads from novel genomes to be reliably identified as homologous. Conserved sequence windows can be used to solve a number of bioinformatic problems in microbial ecology beyond those discussed here, and we plan on exploring these in future. Taken together, SingleM and Sandpiper bring together three sub-fields of microbial ecology—taxonomic profiling, public data analysis and genome-centric metagenomics—in a way that we hope will provide better utilisation of public datasets and improved global context for metagenomic analyses.

## Methods

### Description of the SingleM algorithm

A candidate list of putative single-copy, broad-range marker genes was formed from ribosomal proteins originally derived from PhyloSift(Darling et al. 2014) and GTDB-Tk(Chaumeil et al. 2019) marker gene sets. Some of these genes span both bacterial and archaeal domains, whereas others are restricted to one domain. The set of marker genes was chosen such that each gene is present in either >90% of genomes in Bacteria or >85% of genomes in Archaea, with an average copy number of <1.05. We allowed less than 100% prevalence of these genes in their respective target domains because some reference genomes are incomplete (e.g. MAGs) and some specific lineages have lost certain genes (e.g. Patescibacteria(Méheust et al. 2019)). This heterogeneity is at least partially rescued by robust statistical measures (i.e. trimmed mean) during the ‘condense’ step, detailed below.

The reference database of SingleM (the ’metapackage’) is organised as a collection of ’packages’. Each package details one gene and one window is chosen per gene. To create these packages, Pfam and TIGRfam HMMs associated with each gene were used to extract sequences from GTDB species representatives. The resulting set of SingleM packages were then reduced in number by applying ’singlem pipe’ to the predicted transcripts from GTDB representative species as well as to simulated reads derived from one representative per phyla. Packages that were single-copy in >85% of a domain’s transcripts and simulated reads were included in the metapackage with that domain as a target.

To determine a window position for each gene suitable which is highly conserved and suitable for recruitment of metagenomic reads, raw reads from complex peat metagenomes, which are known to contain reads from a broad range of microbial taxa(Woodcroft et al. 2018) (SRA accessions SRR7151621, SRR7151618 and SRR7151620), were aligned against each HMM using GraftM(Boyd et al. 2018) with parameters ’graftM graft --search_and_align_only’. The generated alignments were then used to identify the position of a 20 amino acid length window containing the greatest number of aligned nucleotides to the marker gene’s HMM, using ’singlem seqs’ with default parameters.

#### Generating marker-wise OTU tables

Raw metagenomic reads are assigned to taxonomically annotated OTUs through the application of several steps, described below. These steps are implemented in the ‘singlem pipe’ subcommand. Many of the parameters detailed can be changed by the user, here only the default parameters are shown. Similarly, a number of performance optimisations are omitted for brevity.

1. The first step in the SingleM algorithm, referred to in the codebase as the ’prefilter step’, is to recruit raw reads to marker gene windows. To hasten this procedure, reads are initially selected by DIAMOND blastx against marker gene sequences from the target domains for each marker. Despite the improved speed afforded by the DIAMOND algorithm(Buchfink et al. 2021), selection of raw reads to align against each marker’s HMM remains the bottleneck in SingleM’s runtime. We take three measures to limit the runtime of this DIAMOND search: (1) use of the DIAMOND ’makeidx’ feature for small reference databases(Edgar et al. 2022), (2) trimming of database sequences to the 20 amino acids in the windows plus 30 amino acids on each side, and (3) sequence dereplication at 60% identity using CD-HIT(Fu et al. 2012) with parameters ‘cd-hit -n 3 -M 0 -c 0.6’. DIAMOND BLASTX is run with parameters ‘diamond blastx --outfmt 6 qseqid full_qseq sseqid --top 1 --evalue 0.01 --block-size 0.5 --target-indexed -c1 --query-gencode 4’. A single database comprising sequences from all markers is used. The output from this step is a set of read identifiers, read sequences and the marker gene they best match to. We found that specifying translation table 4 worked well in practice, because doing so detects those lineages which use translation table 4, but also because inappropriately translated sequences from genomes which use table 11 (the standard bacterial table) were excluded on the basis of sequence dissimilarity. In this default mode, reads are assigned only to their best matching marker gene, which is appropriate for short reads. Long reads, but contrast, may encode genes from multiple markers colocated on a genome. Therefore we suggest the current direct BLASTX approach used by default in SingleM is inappropriate for long reads. If the input to SingleM is a genome (’--genome-fasta-files’), then a quick, rough transcriptome generated by OrfM(Woodcroft et al. 2016) using a minimum gene length of 100 amino acids. Since many of these predicted transcripts are not true genes and may overlap, a dereplication step is applied after marker HMM alignment such that only the longest open reading frame is kept at each locus.
2. Candidate sequences are aligned to the HMMs of their respective marker genes. Translated open reading frames are identified with OrfM using a minimum open reading from size of 24 amino acids (‘orfm -m 72’) and then aligned using the hmmalign tool in the HMMER suite(Eddy 2011).
3. Aligned amino acid sequences are filtered to remove any sequences which do not cover the window. Specifically, any sequences which do not align to both the first and last positions of the window are excluded from further analysis.
4. Sequences are translated from aligned amino acid sequences back into aligned nucleotide sequences, using the matching read sequence. This 60bp nucleotide sequence is then the ‘OTU’ sequence. The redundancy of the genetic code means that these 60bp are a richer source of information than the 20 amino acid sequence when differentiating closely related OTUs and when attempting to apply taxonomic assignment to the species level. This sequence may include gaps, but any inserts are removed so that all OTU sequences are 60bp in length. This consistency of length facilitates efficient comparison of OTU sequences and taxonomic assignment. Sequences containing insertions were also found to be rare in practice.
5. Sequences with the same OTU sequence are aggregated together by exact sequence clustering of the 60bp windows, creating an OTU table. This OTU table can be dereplicated by inexact sequence clustering using the ‘singlem summarise’ subcommand, if desired, though the ‘condense’ algorithm includes correction mechanisms for sequencing error (see below).
6. The ’coverage’ of each OTU is calculated using the established relationship between kmer coverage and read coverage as set out by Velvet(Zerbino and Birney 2008):

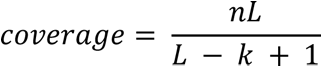 Where n is the number of reads with the OTU sequence, L is the length of the read and k is the length of the OTU sequence including inserts but excluding gaps (usually 60 bp). In practice, each read may have a different length and/or aligned length within the 20 amino acids, so the coverage contribution of each read is calculated separately according to the formula above. The coverage assigned to an OTU is the sum of each read’s contribution.
7. OTUs are assigned taxonomic annotations by matching their nucleotide OTU sequences to a database of species representatives from the GTDB(Parks et al. 2022), using the ’query’ procedure (see below). Sequences are assigned to their closest matching species with a maximum difference of 3bp, since 3 out of 60bp corresponds to 95%, the ANI threshold used for species delineation in the GTDB(Parks et al. 2020). Sequences are matched using the ‘naive’ method of the ‘singlem query’ machinery, described below. When several species have equivalent best hits, the taxonomic assignment of the OTU is then the last common ancestor of these species. The ‘condense’ algorithm incorporates these equal best hits directly to disambiguate taxonomy in these cases (see below).
8. OTUs which are not assigned taxonomy in the previous step are assigned taxonomy via DIAMOND BLASTX. The raw, unaligned and untrimmed read sequences of each OTU are used as input, searching against a database of sequences derived from the OTU’s assigned marker gene. Like the initial read recruitment (prefilter) step, this database consists of protein sequences trimmed to the ∼20 amino acids which align to the HMM window plus 30 amino acids on either side using translation table 4, but unlike the prefilter step the database is not dereplicated. The database also includes protein sequences derived from ’off-target’ species e.g. archaeal sequences from bacterial-only markers. Eukaryotic sequences are also included as off-target, as derived from UniProt truncating the taxonomy to the kingdom level. DIAMOND is run with parameters ‘diamond blastx --outfmt 6 qseqid sseqid bitscore --top 1 –evalue 0.01 --block-size 0.5 --target-indexed -c1 --query-gencode 4’. The taxonomic annotations of these hits are processed in a similar way to the previous step: equal best hits are recorded for later use by ‘condense’. Within an OTU table, the taxonomy of each OTU is calculated by gathering a taxon string for each read in the OTU, which is the last common ancestor of taxons which hit best for each read. Then the taxonomy of the OTU is the most specific taxonomic annotation such that 50% of the reads’ last common ancestors agree. In the generated OTU table and condensed taxonomic profile (see below), no assignment is made to the species level for entries that are assigned taxonomy through DIAMOND BLASTX. Taxonomic annotation made to the genus level at most, since there is insufficient identity on the nucleotide level to be assigned to a specific species. For species where no representative is known to the genus level (novel genera, novel families, etc.), a genus level annotation will be incorrect. In the current implementation, we do not attempt to remedy this and as such interpret genus level assignments as being either correct or representing lineages that are novel at the genus level or higher.
9. The OTU tables generated are optionally output as an ’OTU table’, which is a tab-separated file containing one OTU per line, or an ’archive OTU table’, which is a JSON format file containing more detailed information about each OTU.
10. The OTU table is optionally subject to the ‘condense’ procedure (see below), and output as a ’taxonomic profile’ and/or Krona HTML(Ondov et al. 2011).

#### Query: assigning taxonomy by comparison of OTU window sequences

An OTU window sequence is a 60bp sequence which has been aligned to a marker’s HMM. Unlike a traditional sequence similarity search, which might use a more general local alignment algorithm such as Smith-Waterman to find an optimal alignment between two sequences, comparison between window sequences is a simpler problem. This is because the two sequences are aligned before comparison, since they have both been aligned to the same HMM. Comparing window sequences can therefore be achieved through simple pairwise comparison of the pair of bases at each position in the window.

This simpler problem can further be reframed as a vector similarity search problem, by one-hot binary encoding the base at each position. We represent A with [1,0,0,0,0], T with [0,1,0,0,0], C with [0,0,1,0,0], G with [0,0,0,1,0], and other characters (Ns, gaps or IUPAC codes) with [0,0,0,0,1]. Each position of the 60bp is represented by one of these, and concatenating these across the 60 positions, we generate a binary vector of length 60*5 = 300 for each sequence. For sequences containing only A, T, G and C, the number of positions that differ between two sequences is the Manhattan distance between their vector representations divided by 2. It is divided by 2 since at a mismatching base position, 2 columns will differ.

Calculating these distances can be quickly computed particularly since modern compilers utilise CPU instructions which operate on vectors of bits. If we have one 60bp sequence as a query and a comparatively small number sequences in a database, such as the current number of species in GTDB R214 (85,205 species, each containing ∼1 unique single copy marker gene sequence), then we can compute the most similar set of sequences by brute force, comparing the query sequence against each database sequence. We term this approach the ‘naive’ method.

For larger scale comparison of sequences, the search time can become prohibitive. To speed this search up, the problem can be solved inexactly. The inexact version is known as approximate k-nearest neighbours (approximate kNN), here in 300 dimensional space. Approximate kNN is a well studied problem, particularly since it has many applications in machine learning(Aumüller et al. 2020). However, most implementations assume each dimension is not binary but instead a float value. This likely means that the implementations are not computationally optimised as they might be, but nonetheless provide accurate results. One exception to this is NMSLIB(Boytsov and Naidan 2013), which does provide a binary space implementation. We tested a number of binary and floating point implementations, finding that SCANN(Guo et al. 13--18 Jul 2020) was the most accurate and fast, though ANNOY (https://github.com/spotify/annoy) required less RAM and had a smaller start-up time since it is an on-disk implementation.

Due to the merely approximate results and slightly ill-suited implementations available, we implemented an exact brute force search program, ‘smafa’, and use it as the default window search method (’smafa-naive’ in the SingleM codebase). Implemented in the Rust programming language using needletail (https://github.com/onecodex/needletail), smafa efficiently and exactly finds similar window sequences. It uses the postcard format (https://github.com/jamesmunns/postcard) to store its sequence database with the primary aim of fast database load times. For GTDB 08-RS214, the average marker’s sequences require only a ∼20MB sequence database file.

#### Condense: combining OTU tables from each marker gene into a single taxonomic profile

On their own, the set of OTUs from each marker can be considered a taxonomic profile. However, we provide a method to combine (‘condense’) these into a single taxonomic profile which is advantageous for several reasons. Holistically using the information contained across marker genes is more sensitive, because lower abundance community members may not be represented in each marker’s OTU table. It also allows more specificity in taxonomic annotations, because sequences shared by multiple taxa in one marker’s table may be disambiguated by the sequences observed in another. For instance, if one marker’s OTU table contains a sequence that matches 2 species in one genus (species A and species B), but another marker only contains sequences that match species B, then it is most likely that species B is present in the sample while species A is not. Finally, inspecting one taxonomic profile is simply more convenient than inspecting all 59 individually.

There are some important disadvantages of condensing each markers’ OTU table into a single taxonomic profile, though. In the current implementation, information about the diversity of sequences is not incorporated directly, only their taxonomic affiliation(s) are. For instance, consider a situation where there are two window sequences from different species assigned to a genus G in each of the marker OTU tables, but neither of these species are contained in the reference database (GTDB). The final taxonomic profile will show only coverage of the genus G, with no delineation of lineages at the species level within this genus. In this case, community structure at the species level will not be evident in the condensed taxonomic profile, even though the marker OTU tables show two separate species from the genus are present.

The condense algorithm works in several steps:

1. Any OTUs which have ‘off-target’ taxonomic annotations are removed. These might be Eukaryotic OTUs, or bacterial OTUs which matched archaeal markers, or OTUs not assigned domain-level taxonomy, for instance.
2. Species-wise expectation-maximisation is used to disambiguate the taxonomic affiliation of OTUs that have been assigned to multiple species when matching their nucleotide window sequences to GTDB species nucleotide window sequences. In some cases, window sequences derived from multiple species are identical, and novel strains may map with identical imperfect identity to multiple species. To address these situations, information from other marker gene OTUs is used. Specifically, in this iterative expectation-maximisation procedure, each species is initially assigned equal abundance. Then for each OTU, the coverage is partitioned according to the abundance ratio of species that the OTU matches. The abundance of each species is then re-calculated as the average abundance across the markers (counting only markers targeting the domain to which the species belongs), and the procedure repeated until no species changes in abundance by >0.001 coverage units. A simplified example of this procedure is provided in **Supplementary Note 5**.
3. In order to suppress false positive species that might otherwise be predicted to be present in low abundance by window sequences derived from reads that contain sequencing errors, a ‘shadow abundance’ threshold is applied after calculating the average abundance in the iterative algorithm above. Any species which is present at <10% of its genus’ total abundance, and which is not associated with 10 or more different markers to the exclusion of all other species, is removed. After the expectation maximisation has converged, in rare cases it may still not be possible to disambiguate some sets of species. For these sets, the OTU coverages associated with them are assigned a taxonomy that is the last common ancestor of the species in the set.
4. Genus-wise expectation maximisation is used to disambiguate the taxonomic affiliation of OTUs that have been assigned to multiple taxons through DIAMOND BLASTX. It is unlikely that reads assigned through this method are from species that exist in the reference database since their nucleotide window sequences did not closely match any in the database, so this step seeks only to assign taxonomy down to the genus level, but no further. The procedure is similar to the expectation maximisation used above, except that it assigns taxonomy to genera rather than species. The ‘shadow abundance’ thresholding is also not applied. Coverages from OTUs that have been assigned by nucleotide sequence are included in the calculation of genus-wise coverage, but the taxonomic assignment of these lineages is not modified in the second step.
5. Combination of OTU coverages into a single taxonomic profile. The final profile is created in a step-down approach, where the coverage of each domain is calculated, then the coverage of each phylum, and so on, down to species level. The coverage for each domain is calculated as the trimmed mean of marker-wise coverages, excluding the highest and lowest 10% of values. The coverage of each phylum is then calculated in the same way, but to make it consistent with the domain-wise coverages, the coverage of each phylum in a domain is calculated as a proportion of the overall domain’s coverage. These proportions are the percentage of coverage values assigned either to a phylum (including its taxonomic descendents) or to the domain without further taxonomic specificity, including coverage that has not been assigned to any phylum. This process is then repeated down to species level.
6. The rate of taxonomic assignment to the species level is increased to account for sequencing error. To account for sequencing read error that reduces the level of resolution of an OTU taxonomic assignment from the species to the genus level for known species, 10% of the coverage of each genus is partitioned out to each species, in proportion to their coverage before this step. If <10% of the genus’ coverage is unassigned before application of this step, all of the unassigned coverage is partitioned out instead.
7. The resulting taxonomic profiles are output in a simple tab-separated format and/or KRONA plot(Ondov et al. 2011).

#### Supplement: Adding new genomes to the SingleM reference database

The SingleM ’supplement’ mode takes in a list of genomes in FASTA format, and a reference package (a SingleM ’metapackage’) to be supplemented according to the following procedure:

1. Genomes are filtered for quality using as input a CheckM2(Chklovski et al. 2022) quality file, with the default cutoff of minimum completeness 70% and maximum contamination 10%. This step is optional.
2. Genomes are dereplicated using Galah(Aroney et al. 2024) at 95% average nucleotide identity, so as to only include one representative per species cluster such that the 95%/3bp threshold used in the ’singlem pipe’ is appropriate. Galah is used to choose genomes of highest quality according to the following formula, greedily selecting genomes to include in the supplemented package. The quality formula used to rank genomes is similar to that used in GTDB for species clustering, but only including those scoring criteria that can be calculated from the sequence without homology search. Completeness and contamination values used are those provided in the CheckM2 quality file.

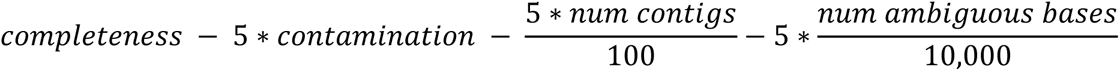
3. Transcripts and protein sequences for each genome are generated using Prodigal(Hyatt et al. 2010). As with GTDB-Tk(Chaumeil et al. 2022), the genome is determined to use the non-standard translation table 4 if both of the following conditions hold, otherwise translation table 11 is used:

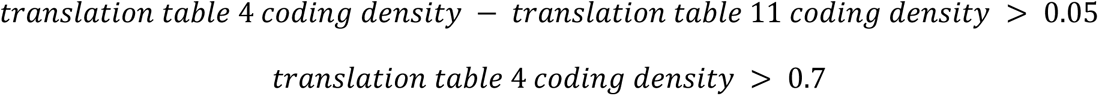
4. Genomes are assigned taxonomy using GTDB-Tk, the database version of which must be equal to that used to generate the original metapackage. Genomes which are assigned a species level taxonomy are excluded since they do not add new species.
5. Protein sequences from remaining genomes are searched with HMMSEARCH using the HMMs of each SingleM marker gene with a default e-value of 1e-20. Each protein is assigned to at most one marker gene.
6. SingleM ’pipe’ is run on the transcripts of hit proteins to gather 60bp sequences for use with ’smafa-naive’.
7. Further bookkeeping procedures are carried out and a final supplemented metapackage output.

### Reduced genome marker searching

To determine the number of markers contained within extremely reduced bacterial genomes (**Supplementary Data 1**), SingleM ’pipe’ was run using default parameters with the genome sequence as input, outputting an OTU table. The number of markers was the number of unique markers which remained after removing ’off-target’ markers (i.e. archaeal markers which are not in the bacterial set, but may nonetheless be encoded in some bacteria) using SingleM ’summarise --exclude-off-target-hits’. We note that many of the tested genomes use translation table 4, but we report the number of markers found by SingleM, which currently assumes translation table 11 during ’pipe’ mode.

### Benchmarking

Benchmarking was carried out within Snakemake(Köster and Rahmann 2012) pipelines, which are available at https://github.com/wwood/singlem-benchmarking.

#### Novel lineage detection

To benchmark detection of novel lineages, a pipeline was created which simulated read sequences which were from lineages present in GTDB R214 but not GTDB R207. Specifically, 120 genomes were chosen where the GTDB R214 taxonomy contained no species representatives that were in GTDB R207 (regardless of their assigned taxonomy). At each level of novelty (from species to phylum), 20 of the highest quality genomes (calculated as CheckM1(Parks et al. 2015) completeness - 5 x contamination) were chosen, with as close to 10 Archaea as possible. The chosen genomes were sometimes from the same novel lineage. To enable direct comparison with profiling tools such as Bracken which estimate the number of reads from each lineage, rather than the relative abundance of each lineage(Sun et al. 2021), the known and novel genomes were chosen to have genome sizes as similar as possible.

To run each benchmark, reads were simulated from 120 communities each containing a novel genome and a known genome (either *Staphylococcus aureus* assembly GCF_001027105.1 or *Methanobrevibacter ruminantium* assembly GCF_000024185.1), at equal read coverage of 10X. Paired-end 150bp reads were simulated using ART version 2.5.8(Huang et al. 2012) with parameters ’-ss HSXt -p -l 150 -f 10 -m 400 -s 10’. To test against the gold standard, the output of each tool was first converted to the ’condensed profile’ format, the default SingleM taxonomic profile output format using custom scripts available in the benchmarking codebase, and then further converted to biobox format(Belmann et al. 2015) and compared to gold standards using OPAL(Meyer et al. 2019) v1.0.11. To test detection (**Figure 2**), communities were compared at the kingdom level. To benchmark classification of novel lineages lower ranks were used (excepting Kaiju and MAP2B for which no GTDB R207 reference database was available). Reference databases were transferred to local scratch space to minimise the effect of IO wait on runtimes.

SingleM ’pipe’ v0.15.0 was run with default parameters. MetaPhlAn v4.0.6 was run by first concatenating paired-end reads into a single gzip compressed FASTQ format. Taxonomy assignments were converted to GTDB using mpa_vOct22_CHOCOPhlAnSGB_202212.pkl with the supplied sgb_to_gtdb_profile.py script. mOTUs v3.1.0 ’profile’ was run using default parameters and converted to condensed format using the provided ’mOTUs_3.0.0_GTDB_tax.tsv’ mapping file. Sourmash 4.8.2 was run using the GTDB 07-RS207 reference database using ’sourmash sketch dna -p k=21,k=31,k=51,scaled=1000,abund’, and using the median_abund as the abundance measure. The Kraken2+Bracken workflow used the GTDB database built by Struo2(Youngblut and Ley 2021). Kraken2 v2.1.2(Wood et al. 2019) was used with ’kraken2 –report .. –paired ..’ followed by Braken git commit 88b7738 using ’-t 10’ and ’-l’ for each taxonomic level. This produced a report for each taxonomic level, which was then converted to condensed format. To compare classification accuracy, the taxonomic annotation of the novel genome in GTDB 07-R207 was estimated using GTDB-Tk(Chaumeil et al. 2022) version v2.1.0.

The taxonomy assignments of Kaiju and MAP2B are not based on GTDB R207 taxonomy, so these tools could not be fully benchmarked against the rest of the tools. To assess their ability to detect novel lineages, we converted taxonomy assignments to the kingdom level (i.e. Bacteria or Archaea) and compared them on this level only. Kaiju 1.9.2 was run using the progenomes 2021-03-02 database, as we are unaware of any GTDB-based reference database. Paired-end reads were concatenated together and provided to the ’kaiju’ executable followed by ’kaiju2table -r phylum’. Kingdom level taxonomies were derived using pytaxonkit(Shen and Ren 2021) (https://github.com/bioforensics/pytaxonkit). MAP2b(Sun et al. 2023) v1.5 was run using the data specified in its ‘config/GTDB.CjePI.database.list’ file, a database generated from GTDB R202.

#### Profiling of communities of known species

To benchmark profiling tools against communities of species present in the reference database, a similar set of procedures and reference databases were used. Reads were simulated according to the abundance profiles in the 10 CAMI 2(Meyer et al. 2022) ’marine’ communities. All entries in the coverage definition file (‘OTU’ or otherwise) were simulated as microbial genomes, for an average of 469 simulated genomes per sample. To emulate a more realistic community, genomes which were not species representatives were chosen for simulation. To reduce bias in the chosen species towards highly sequenced species, for each species, only those genomes in the top 20 genomes ordered by completeness - 5*contamination were included in the set to choose from. Genomes were chosen at random from the remaining set of genomes to include in the profiling benchmark. Runtime and RAM usage stats were collected using the ‘s’ and ‘max_rss’ columns output by the Snakemake benchmark directive. Figures were generated using R(Ihaka and Gentleman 1996), ggplot2(Wickham 2016) and patchwork(Pedersen 2014).

### Generation of Sandpiper dataset

A set of metagenomes to be analysed were collected according to the following criteria, querying Google BigQuery via SQL where each of the following conditions was true: (1) The ’librarysource’ was ’metagenomic’, or the ’organism’ was a descendent of the ’metagenome’ taxonomy, (2) The ’platform’ was ’ILLUMINA’, (3) ’consent’ was ’public’, (4) ’mbases’ was >1000 or ’libraryselection’ was ’RANDOM’ and mbases was > 100, (5) mbases was <= 200,000, (6) librarysource was not ’VIRAL RNA’ or ’METATRANSCRIPTOMIC’ or ’TRANSCRIPTOMIC’.

Metagenomes were analysed using kubernetes on Google GCP or Amazon AWS. Metagenomes were copied from AWS in .sra format and streamed to SingleM ’pipe’ using Kingfisher(Woodcroft et al. 2024). The git commit of SingleM used was e97d171 and the reference database used was ’S3.metapackage_20211101.smpkg’ (DOI 10.5281/zenodo.5739612), based on GTDB 06-RS202. We note that this version of SingleM did not specify ’--query-gencode 4’ in its initial DIAMOND BLASTX, as the current version does, so lineages which use translation table 4 are likely underrepresented in these profiles. Outputs were generated in ’archive OTU table’ format and later processed using ’singlem renew’ to update the taxonomy annotations of each genome to GTDB R214 version (DOI 10.5281/zenodo.7955518) using SingleM v0.16.0. Taxonomic profiles are available at DOI 10.5281/zenodo.10547494.

The Sandpiper website was built using Flask (https://flask.palletsprojects.com) and Vue (https://vuejs.org/). The source code is available at https://github.com/wwood/sandpiper/ and incorporates a list of manually curated corrections to NCBI-derived project and sample metadata available at https://github.com/wwood/public_sequencing_metadata_corrections.

#### Biome-wise breakdowns of taxonomic profiles

The biome each metagenome was derived from was mostly derived from the ’organism’ field stored in the biosample associated with each metagenome at NCBI. However, given the large number of metagenomes assigned to an undifferentiated organism ’metagenome’, we trained a machine learning classifier to predict whether a metagenome is either eukaryotic host-associated or ecological based upon its taxonomic profile. Using metagenomes annotated as ’organismal metagenomes’ as host-associated and ’ecological metagenomes’ as ecological as the gold standard, an XGBoost(Chen and Guestrin 2016) model was trained, using five-fold cross validation. To minimise overtraining, we grouped metagenomes by their BioProject such that metagenomes from one BioProject were never included in both the training and test sets at the same time, using the GroupKFold function of sci-kit learn(Pedregosa et al.). Taxonomic profiles were input using the relative abundance of phylum, class or orders.

Models trained at each of these taxonomic levels showed similar performance during cross-validation (∼93% accuracy). The final predictor was trained on all of the gold standard data with order-level taxonomic profiles as input. When a metagenome was assigned an organism which is eukaryotic host associated or ecological in its metadata, that annotation was used for analysis here and on the Sandpiper website. Biomes more specific (e.g. soil metagenome) were taken directly from biosample metadata. The predictor is made available at https://github.com/wwood/singlem_host_or_ecological_predictor.

#### Fractions of metagenomes assigned to the species level

To establish the fractions of available communities classified at the species level at the current time, the default GTDB R214-based SingleM reference database (metapackage) was supplemented with genomes from the ‘UHGG’ version 2(Almeida et al. 2021), ’SPIRE’ (excluding “specI” isolate genomes)(Schmidt et al. 2023), ’SMAG’(Ma et al. 2023), ’GEM’(Nayfach et al. 2021) MAG collections, as well as those from derived from Oceans by Paoli et. al.(Paoli et al. 2022). SPIRE species representative MAGs were downloaded from https://spire.embl.de/downloads, SMAG from https://zenodo.org/records/8223844, GEM from https://portal.nersc.gov/GEM/genomes/fna, and Ocean MAGs from https://sunagawalab.ethz.ch/share/microbiomics/ocean/suppl_data/representative-genomes-fasta.tar.gz. All genomes were quality controlled using CheckM2 v1.0.2(Chklovski et al. 2022), assigned taxonomy using GTDB-Tk v2.3.0(Chaumeil et al. 2022) ‘classify_wf’. Any genomes <50% complete, >10% contaminated or assigned to a species level taxonomy by GTDB-Tk were excluded. Genes were called using “prodigal-runner” to run prodigal choosing translation table 4 or 11 as appropriate (https://github.com/wwood/prodigal-runner git commit c5f7713) based on the process established by GTDB-Tk(Chaumeil et al. 2022). The total set of MAGs was dereplicated at 95% ANI using Galah(Aroney et al. 2024) git commit f199654 which used skani(Shaw and Yu 2023). These data were input into “singlem supplement” to generate a new metapackage, which is available at DOI 10.5281/zenodo.10360136. The profiles generated are available at DOI 10.5281/zenodo.10547501.

This new metapackage was used with ‘singlem renew’ to reannotate the taxonomy of OTU sequences in SRA metagenomes, and to regenerate condensed profiles. We note that while this approach was used to provide an estimation of the known species fraction inclusive of these MAG data, and for high level taxonomic overviews, it is unsuitable for general purpose community profiling because taxonomic assignment of genomes was made without proper estimation of the taxonomic structure between the species level and the highest level of taxonomy provided by GTDB-Tk. As a concrete example, if two novel species are assigned to the same taxonomic family (and not to any genus), then ‘singlem supplement’ currently assumes they are from distinct genera, even if they are actually congeneric.

The known species fraction for each metagenome was calculated simply as the sum of coverage values reported in the SingleM profile divided by the total of coverages assigned to all taxonomic levels. To address potential biases arising from metagenomes with limited sequencing depth, reported mean and median values are amongst those metagenomes with >50 total coverage in the SingleM taxonomic profile and total sequence depth >1 Gbp. Biome-wise breakdown of known species fractions and phylum-wise relative abundance (**Supplementary Data 3**) were taken from the NCBI ‘organism’ metadata entry. Human samples were those with ‘human’ as a substring of their organism entry, or had organism ‘gut metagenome’, ‘feces metagenome’ or ‘oral metagenome’. Mouse, pig, bovine metagenomes were found by searching for organisms containing each as a substring. Marine samples were those with ’seawater metagenome’ or ’marine metagenome’ as their organism. Plant, soil, sediment, freshwater and aquatic metagenomes were identified based on exact matching of their organism e.g. “plant metagenome” to identify plant metagenomes.

The default GTDB R214 SingleM metapackage was used for the following analyses. To ascertain the fraction of available communities classified at the species level over time, the NCBI datasets tool (https://github.com/ncbi/datasets) was used to download the genome summary in JSON format for each species (whether a species representative or not) in GTDB R214, and the submission date for each genome found using jq -rc ’.reports[] | [.accession,.assembly_info.submission_date] |@tsv’.(https://jqlang.github.io/jq/). The earliest submitted genome from each GTDB species was then calculated as the first year in which any genome in the species cluster was submitted. The set of metagenomes included in the analysis also had to pass these criteria: (1) The total sample coverage had to be >50 to ensure adequate microbial sequencing depth, (2) the coverage assigned to any one genus could not exceed 90% of the total coverage to exclude single cell genomes. The date of the metagenome was the ’releasedate’ in the metadata, collected using ’kingfisher annotate’(Woodcroft et al. 2024). To determine the fractions of metagenomes which not only have genomic representation but are also present in isolate culture collections, the GTDB auxiliary file ’hq_mimag_genomes_r214.tsv’ (https://data.gtdb.ecogenomic.org/releases/release214/214.0/auxillary_files/) was used to gather a list of GTDB species representatives that are known to be isolated.

### Targeted genome recovery

For genome recovery targeted at Muirbacteria, Wallbacteria, Riflebacteria and Fusobacteria, the set of samples which contained coverage of each of these phyla was extracted from Sandpiper, when it was annotated with GTDB R207. For each of these samples, the total coverage of taxons which were (1) assigned a taxonomy to one of the target phyla and (2) not assigned to the species level (the ’non-species’ coverage) was tabulated for each phyla. The set of chosen samples for targeted genome recovery were those which had a high non-species coverage (>10X coverage) and high ratio of non-species coverage to coverage assigned to the species level in the phyla (>90%). Corresponding metagenomic data was downloaded with Kingfisher(Woodcroft et al. 2024). MAGs were recovered with Aviary (git commit da0efd0)(Creators Newell, Rhys J. P. Aroney, Samuel T. N. Zaugg, Julian Sternes, Peter Tyson, Gene W. Woodcroft, Ben J.), assembling with metaSPADES(Nurk et al. 2017) and binning with CONCOCT(Alneberg et al. 2014), MaxBin2(Wu et al. 2016), MetaBAT(Kang et al. 2015, 2019), SemiBin(Pan et al. 2022) and VAMB(Nissen et al. 2021). Bins from each were combined using DAS Tool(Sieber et al. 2018). Some samples were manually assembled outside of Aviary using megahit v1.2.9(Li et al. 2015) since metaSPAdes(Nurk et al. 2017) (the Aviary default) cannot use single-ended metagenomic data as input. Only one metagenome was used to inform binning via differential coverage, the metagenome used for assembly. Genome quality was assessed with CheckM2(Chklovski et al. 2022). The reported success rate (87%) is only amongst those metagenomes where the assembly and binning steps successfully finished (**Supplementary Data 4**).

## Data availability

SingleM reference databases corresponding to GTDB R207 and R214 are available at DOI 10.5281/zenodo.7582579 and 10.5281/zenodo.7955518 respectively. The reference database used for the initial screen of public metagenomes is available at DOI 10.5281/zenodo.5739612 and the reference database supplemented with genomes not yet in GTDB is available at DOI 10.5281/zenodo.10360136. GTDB-based profiles of public metagenomes are available at DOI 10.5281/zenodo.10547494, and reference-supplemented profiles at 10.5281/zenodo.10547501. Metagenome-assembled genomes from Muirbacteria, Wallbacteria, Riflebacteria and Fusobacteria have been deposited at Zenodo under DOI 10.5281/zenodo.10162715.

## Code availability

SingleM, sandpiper and smafa software are made available under a free software licence at https://github.com/wwood/singlem, https://github.com/wwood/sandpiper/ and https://github.com/wwood/smafa, respectively. SingleM and smafa are available through BioConda (https://anaconda.org/bioconda/singlem), and distributed through PyPI (https://pypi.org/project/singlem/) and crates.io (https://crates.io/crates/smafa) respectively. SingleM is also available through DockerHub (https://hub.docker.com/r/wwood/singlem). Workflows used for benchmarking are available at https://github.com/wwood/singlem-benchmarking and the predictor of sample eukaryotic host-association at https://github.com/wwood/singlem_host_or_ecological_predictor.

## Acknowledgements

B.J.W. was supported by Australian Research Council grants (#DE160100248, #DP230101171, #FT210100521). The project was supported by the EMERGE National Science Foundation (NSF) Biology Integration Institute (#2022070) and Genomic Science Program of the United States Department of Energy (DOE) Office of Biological and Environmental Research (BER), grants DE-SC0004632, DE-SC0010580 and DE-SC0016440. Cloud computing was generously contributed by Amazon Web Services (AWS) and Google Cloud (GCP). We thank the many who have provided helpful comments including Timothy Lamberton, Pierre-Alain Chauneil, Donovan Parks, David Wood, Phil Hugenholtz, Raphael Eisenhofer, Christian Rinke, Jiarul Sun, Apoorva Prabhu, Julian Zaugg, Steve Robbins, Andy Leu, Virginia Rich, Luis Pedro Coelho, Yu Yang, Caitlin Singleton and beta testers of the software. We thank high performance computing system administrators Brian Kemish, Chris Williams and Hamish McDonald. We gratefully acknowledge the large collective effort expended in administering, gathering, sequencing and uploading metagenomic datasets into the public domain.

## Author Information

### Contributions

B.J.W., S.T.N.A., and R.Z. developed the SingleM algorithm, in part under the supervision of G.W.T. B.J.W. and M.C. applied it to public datasets under the supervision of L.B. B.J.W., S.T.N.A. and J.A.M.M. analysed the Sandpiper data. B.J.W. and J.A.M.M. developed the host-association machine learning algorithm. B.J.W., S.T.N.A., R.Z. and J.A.M.M. wrote the manuscript with input from G.W.T. All authors reviewed and approved the final version of the manuscript.

