## Supplementary Notes Figures and Tables for "SingleM and Sandpiper: Robust microbial taxonomic profiles from metagenomic data"

### Supplementary information, figures and table for “SingleM and Sandpiper: Robust microbial taxonomic profiles from metagenomic data”

#### Supplementary Note 1

##### Species level classification using a 96.7% identity threshold within conserved windows

We compared the use of different thresholds to assign a species-level taxonomy to OTU sequences. If the similarity of the OTU sequence is greater than this threshold, then a genus-level taxonomic assignment is instead made, using DIAMOND BLASTX (**Figure 1**, methods). We tested 57bp out of 60, which is a natural choice since it corresponds to the ANI threshold used by GTDB to delineate species. We also tested the next most stringent identity of 58bp out of 60 (corresponding to 96.7% identity). We reasoned that a more stringent threshold might be more appropriate since OTU sequences are derived from highly conserved stretches of highly conserved genes, and so may diverge at a slower rate than entire genomes.

To test these thresholds (and others not shown), we chose genomes from genera with at least three species that had at least three genomes in GTDB R214. Taxonomic profiles from three genomes per genus were generated, assigning taxonomy using the two thresholds in two different ways. First, the default database was used to test how many species-level assignments are made when they *should* be. Secondly, taxonomy was assigned using a database where the species being studied was excluded, to test how many species-level assignments are made when they *should not* be.

These tests were carried out by running SingleM ‘pipe’ with an R214-based reference database (version 3.2.1), assigning taxonomy to the genus level using ‘--assignment-method diamond’ on each genome’s transcripts to generate ‘archive OTU’ tables. These tables were modified by changing their taxonomic annotations to be species-specific, before passing them to ‘singlem condense’ mode to generate taxonomic profiles.

The 57bp cutoff provided more correct species-level assignments when the correct genome was present than the 58bp cutoff (93.8% vs 93.1%), but using a 58bp cutoff reduced incorrect species-level assignments when the correct genome was excluded (from 25.8% to 19.7% false positives, **Supplementary Table 2**). Minimal differences were observed between the cutoffs in the benchmark of known species (<0.5% difference in Bray-Curtis dissimilarity to the gold standard). On balance, we decided to implement the 58bp cutoff as the default since it produced fewer false positive species identifications when the correct species was missing from the reference database.

We emphasise that the 19.7% rate of false positives observed for the final 58bp threshold does not translate into an estimation that 19.7% of species-assignments in community profiles generated by SingleM from arbitrary metagenomes are false, because (1) many genera in GTDB do not contain 3 or more species, and (2) each OTU sequence may or may not be contained in the reference database. It is unclear exactly what the rate of false positives is in arbitrary metagenomes, but the rate will likely be substantially less than 19.7% because correct assignment will reduce the fraction of incorrect species assignment amongst the total of all species assignments. The rate of false positive species identification is discussed further in **Supplementary Note 4**.

#### Supplementary Note 2

##### Profiling accuracy on the known species benchmark

The known species benchmark was further interrogated to assess specific aspects of each tools' performance. For most species present in the benchmark at high true coverage, the coverage predicted by SingleM was observed to be similar (**Supplementary Figure 2**). However, a minority of these species were not predicted to be present at all. In these cases it was observed that strain present in the mock community was comparatively distinct from its GTDB genome representative i.e. close to the 95% species ANI boundary. In these cases, while not predicting the species to be present, the genus was predicted to be present. Further evidence for this was observed when plotting the predicted versus actual relative abundance of individual genera, where the false negative rate was markedly reduced (**Supplementary Figure 3**), which is reflected in a more accurate overall prediction of the community profile (**Supplementary Figure 4**).

Marker based community profiling tools such SingleM are limited in their ability to detect and precisely estimate the relative abundance of species present at low abundance, since they restrict analysis to small sections of each genome. Beyond the global threshold defined as part of the SingleM algorithm, where species predicted to be present at <0.35X coverage are removed by design to reduce noise, an increased error rate was observed for lower abundance members within the known species benchmark (**Supplementary Figure 5**). A full set of standard benchmarking statistics is shown in **Supplementary Data 2**.

#### Supplementary Note 3

##### Ability of SingleM to detect symbionts with highly reduced genomes

The limits of SingleM's marker based approach to detect novel lineages were further tested by applying it to six highly reduced symbiont genomes (<300kb). Although these genomes contained fewer marker genes than average, each encoded at least 19 marker genes (i.e. at least 54% of the bacterial marker set) (average showing these lineages remain detectable despite their extensive gene loss (**Supplementary Data 1**). However, while they are detectable, their abundance will be underestimated by the current implementation of SingleM since a robust average is taken across the marker set to estimate relative abundance of each lineage (see methods).

#### Supplementary Note 4

##### Overclassification rates at the species level

We investigated the extent to which SingleM and other community profiling tools incorrectly assign relative abundance to species when the correct species is not present in the reference database, which we refer to as "overclassification". For each level of novelty, the generated profiles at each level of taxonomic novelty were inspected, tabulating the fraction of the community which was incorrectly assigned to the species level (**Supplementary Figure 6**). Excepting the simulated known genome, a perfect tool would assign 0% of the community to the species level, since the novel species are not present in the reference database. In this benchmark, little to no overclassification was observed in communities containing novel phyla, classes, orders, families or genera by SingleM or other tools tested (MetaPhlAn, mOTUs, and sourmash) with the exception of Kraken2+Bracken, which averaged 14% overclassification (s.d. 13%). However, for genomes novel at the species level, all tools exhibited some level of overclassification. SingleM overclassified by an average of 16% (s.d. 17%) these genomes (median 9%), a rate which compared favourably with all other tools tested (**Supplementary Figure 6**). This erroneous behaviour was often observed in situations where the novel genome was comparatively similar to those already in the database. For instance, each of the top 3 genomes which were overclassified the most (i.e. the three novel genomes which had the highest coverage erroneously assigned to the species level) had 94% average nucleotide identity to the genome they were misclassified as, only slightly more divergent than the 95% threshold used by GTDB for defining species.

Conversely, using the "marine" benchmarks, which were composed entirely of known species, we tabulated how much of each simulated community was not assigned a species-level taxonomy i.e. "underclassification". A perfect tool would assign 100% of the community to the species level since all species are known. SingleM assigned an average of 87% (s.d. 2.4%) of the community to the species level in these communities.

#### Supplementary Note 5

##### Worked expectation-maximisation example

A simplified example of the species-level expectation maximisation procedure is detailed below, imagining a scenario with 2 OTUs. One of the OTUs (OTU1) has an ambiguous taxonomy assignment, and might be either of 2 species, while the other OTU (OTU2) is uniquely assigned to one of these species (species 2). Both species are targeted by both markers.

1) **First we set initial conditions:**

| | |
|---|---|
| Coverage of species 1 | 1 |
| Coverage of species 2 | 1 |

2) **Partition coverages of OTUs**

| OTU | Coverage calculation | Coverage |
|---|---|---|
| OTU1 (species 1) | $1 \times \frac{10}{1+1}$ | 5 |
| OTU1 (species 2) | $1 \times \frac{10}{1+1}$ | 5 |
| OTU2 (species 2) | $1 \times \frac{8}{1}$ | 8 |

3) **Re-estimate species-wise coverages**

We estimate the total coverage as the sum of coverages from each OTU, divided by the number of markers targeted to it. For this simplified example, both species are targeted by the 2 markers (and no others), so we divide by 2.

| | Coverage calculation | Coverage |
|---|---|---|
| Species 1 | $\frac{5}{2}$ | 2.5 |
| Species 2 | $\frac{5+8}{2}$ | 6.5 |

At least one species has changed in coverage by > 0.001, so we iterate again.

4) **Re-partition coverages of OTUs, repeating step (2)**

| OTU | Coverage calculation | Coverage |
|---|---|---|
| OTU1 (species 1) | $2.5 \times \frac{10}{2.5+6.5}$ | 2.8 |
| OTU1 (species 2) | $6.5 \times \frac{10}{2.5+6.5}$ | 7.2 |
| OTU2 (species 2) | $2.5 \times \frac{8}{2.5}$ | 8 |

5) **Re-estimate species-wise coverages, repeating step (3)**

| | Coverage calculation | Coverage |
|---|---|---|

| | | |
|---|---|---|
| Species 1 | $\frac{2.8}{2}$ | 1.4 |
| Species 2 | $\frac{7.2 + 8}{2}$ | 7.6 |

At least one species has changed in coverage by > 0.001, so we iterate again. The process ultimately ends up with all the coverage from OTU1 being assigned to species 2, and species 1 is assigned zero coverage since it falls below the global coverage cutoff of 0.35.

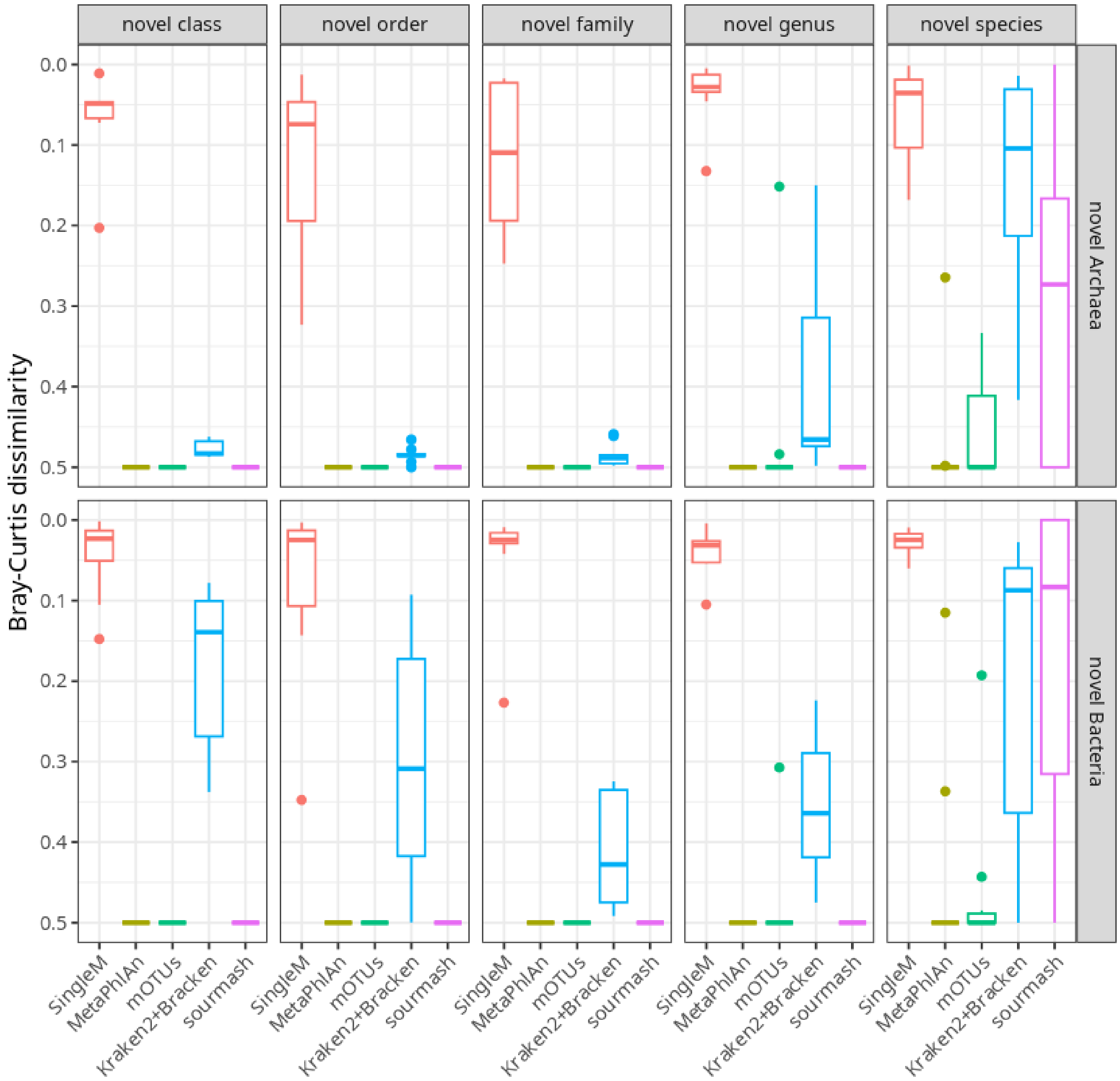


**Supplementary Figure 1: Classification accuracy on novel lineages at the second highest level of possible precision**. Using the novel lineage benchmark data, genomes which were new in GTDB R214 were assigned R207 taxonomy using GTDB-tk, and the two community profiles were compared at the second highest possible level of taxonomic resolution given the constraints of R207 taxonomy i.e. at kingdom level for novel classes, phylum level for novel orders, class level for novel families, order level for novel genera and family level for novel species. The classification accuracy observed for SingleM here was nearly as high as its accuracy in the detection task.

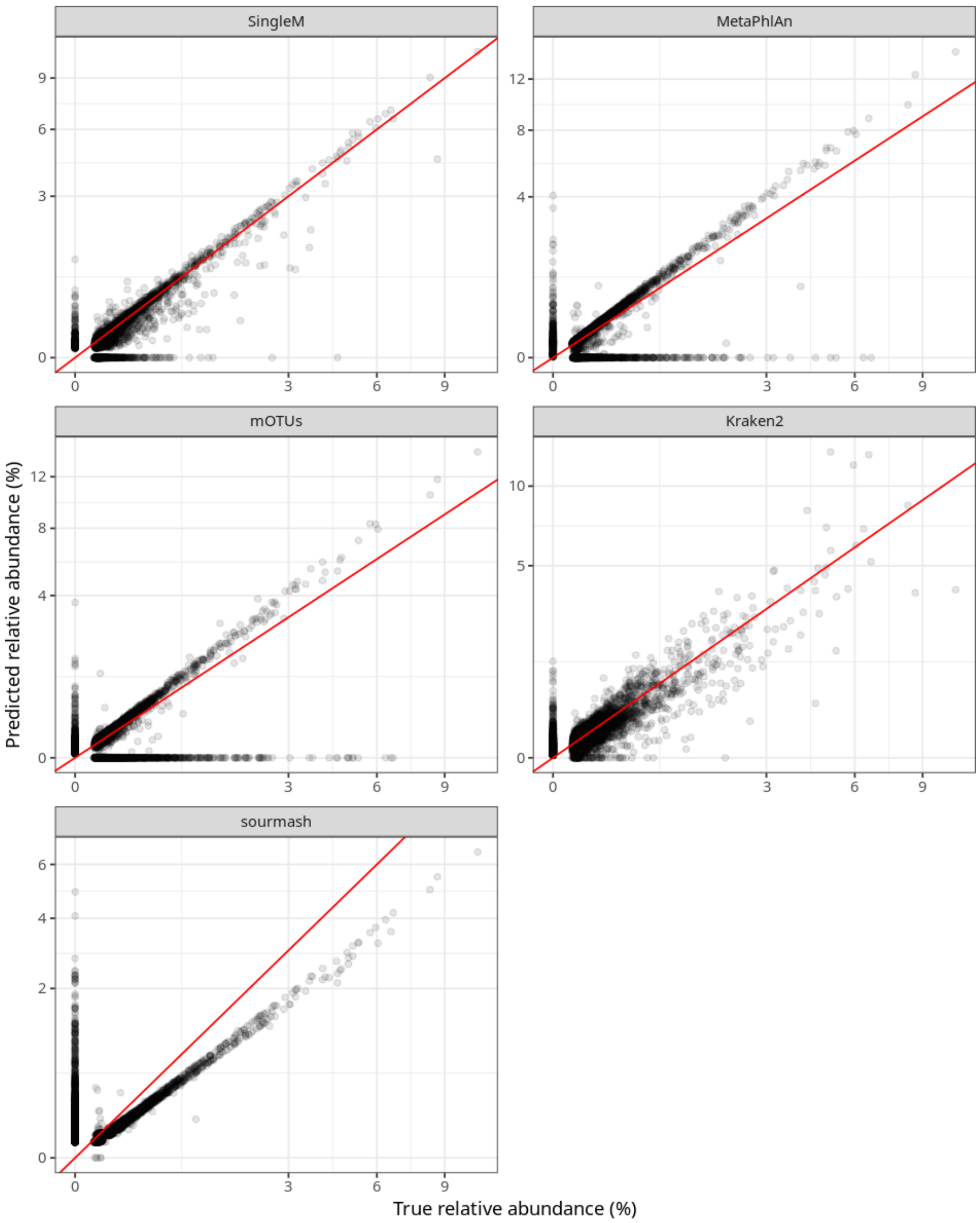


**Supplementary Figure 2. Simulated and predicted relative abundances of individual species on the communities derived from the known species benchmarks.** Each dot represents a GTDB species, plotted by the true and predicted relative abundance for each tool. All species either predicted or simulated to be present from all 10 known species benchmarks are plotted with both axes in square root scale, with each panel's axis limits differing slightly. The red line shown depicts where the predicted relative abundance equals the true relative abundance, so points near that line indicate good performance of the tool for that species. Points along the x-axis indicate false negatives i.e. the species in the mock but predicted absent by the tool, and points along the y-axis indicate false positives i.e. predicted present by the tool when the species was not present in the mock.

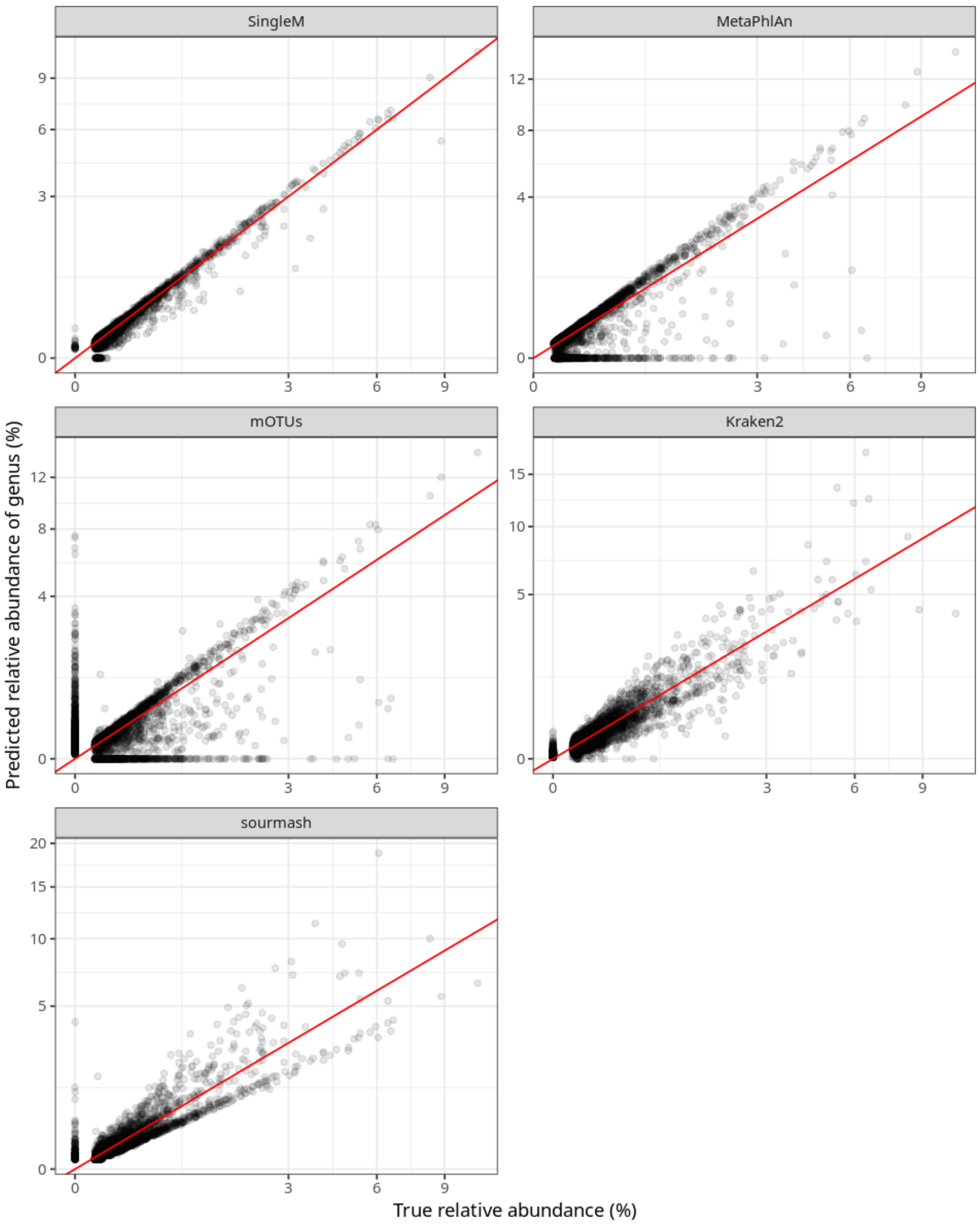


**Supplementary Figure 3. Simulated and predicted relative abundances of individual genera on the communities derived from the known species benchmarks.** This plot is similar to **Supplementary Figure 3**, however instead of individual species being plotted, each point represents the sum of relative abundances of an individual genera. Compared to the species level, most tools have a much reduced false positive rate. The false negative rate for SingleM is also markedly reduced.

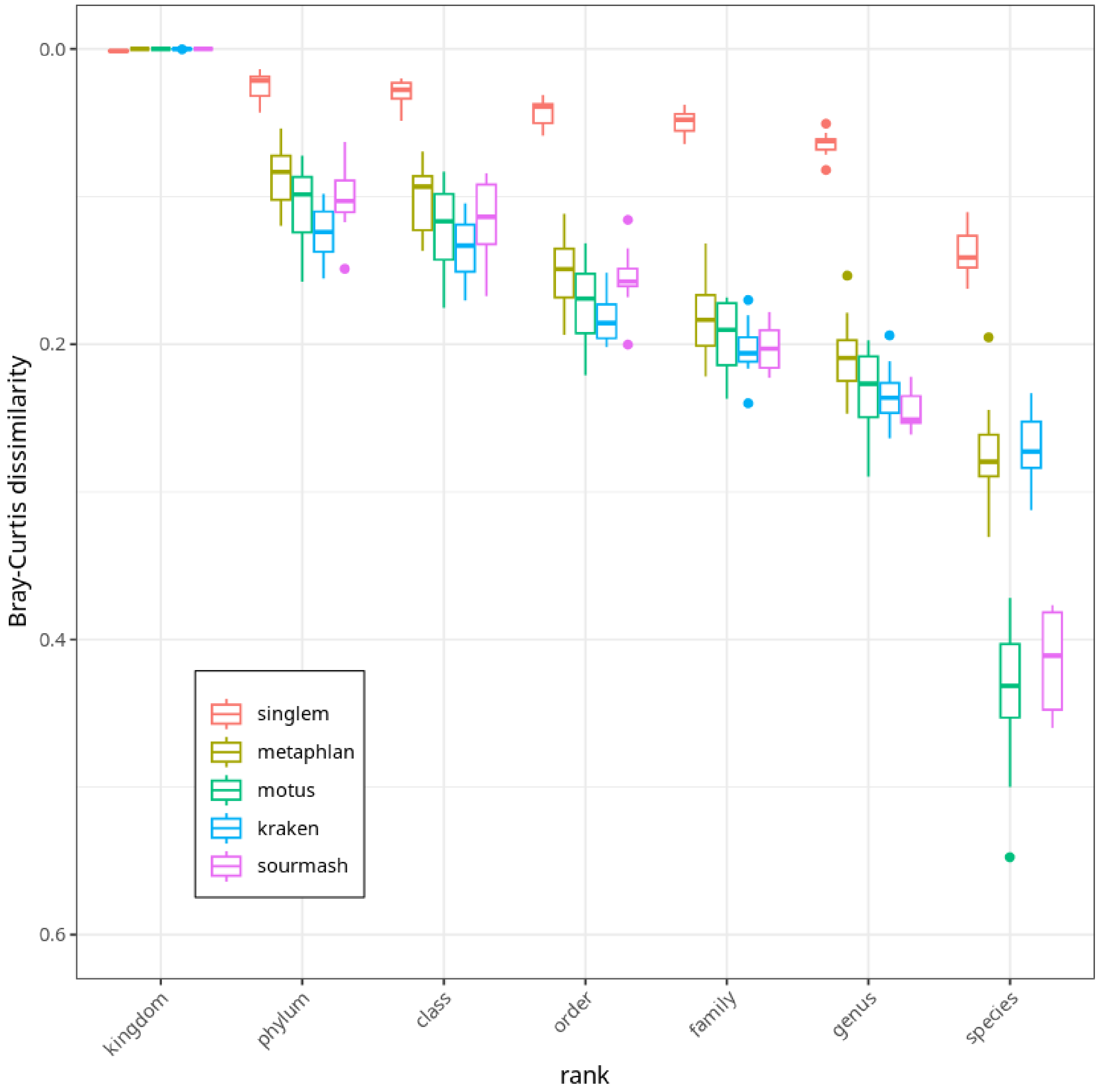


**Supplementary Figure 4. Overall benchmark accuracy on known species benchmarks at each taxonomic level.** For each of the 10 communities simulated as part of the known species benchmark, the Bray-Curtis dissimilarity of the predicted community profile to the true community profile is plotted, at each level of taxonomy, with the boxplots depicting the 10 individual results. SingleM and CoverM performed best in this benchmark.

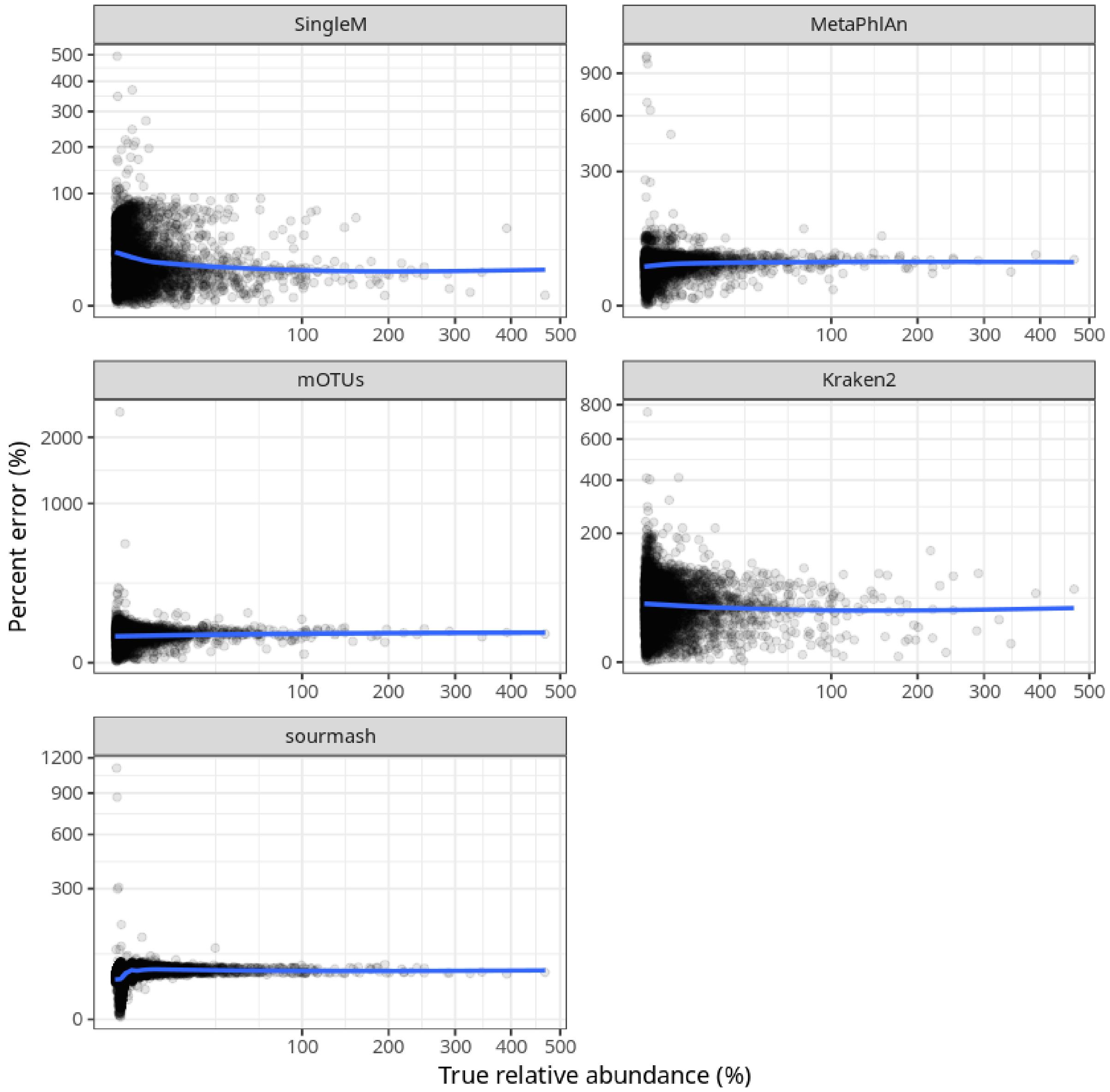


**Supplementary Figure 5. Error rates on the known species benchmark versus true relative abundance**. Each point plotted represents a species in the 10 communities simulated, with false positive and false negative species excluded. Only points where the tool and the true abundance is >0 are plotted. The scale of the y-axis (error rate) differs between panels. The error rate ([true coverage - predicted coverage]/true coverage * 100%) is flat (but variable) for most tools except SingleM, where an increased error rate is observed for species present at lower abundance.

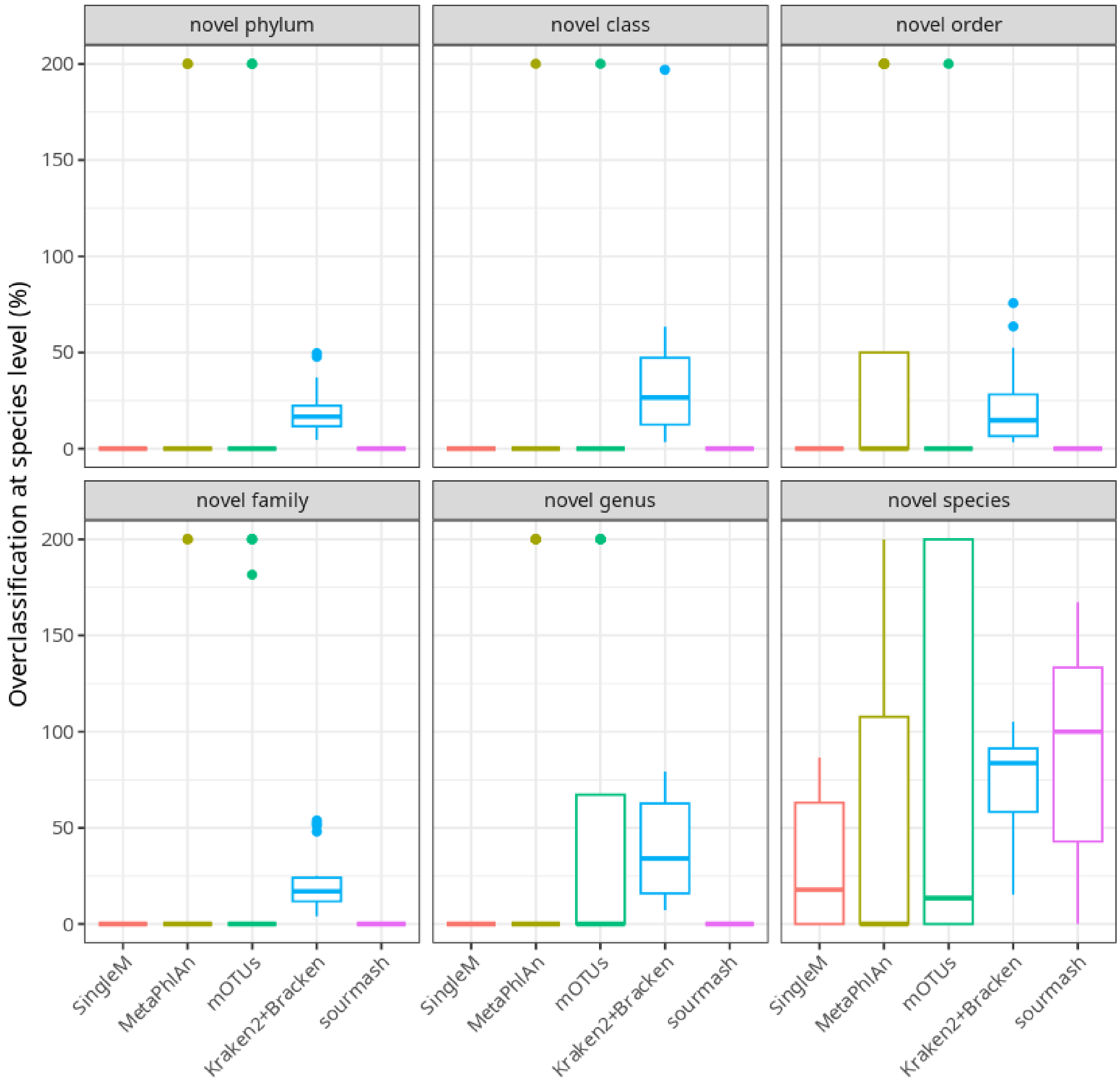


**Supplementary Figure 6: Rates of overclassification to the species level in the novel species benchmark**. The fraction of the community which was incorrectly assigned to the species level was calculated relative to the relative abundance of the known species in each mock community. Some percentages for some tools are greater than 100% owing to misclassification of the known species' reads as well as the novel species' reads.

#### Supplementary Table 1.

Relative abundance of genomically sequenced and cultured species in metagenomes from the top 5 categories within the "organismal metagenome" (i.e. eukaryotic host-associated) and "ecological metagenome" classes recorded in Sandpiper. To address potential biases arising from metagenomes with limited sequencing depth, metagenomes were filtered to only include those with >50 total coverage in the SingleM taxonomic profile and total sequence depth >1 Gbp. Metagenomes are assigned to classes and biomes according to their annotated NCBI "organism" taxonomy. "Subtotal" for the *Host* class include all unlisted biomes predicted to be host-associated (see methods). "Subtotal" for the *Ecological* class includes only marine, soil, sediment, freshwater, and aquatic samples; as many biomes within this class are sourced from "man-made" environments (e.g. wastewater, food, bioreactor, etc), which tend to have higher known/cultured species fractions (see Wastewater) and skew the summary statistics presented here. The known species fraction in human samples was slightly reduced using the supplemented reference database despite this database including many more genomes. This was likely due to challenges assigning taxonomy within genera where large numbers of species are represented.

| | | ***Known species fraction (GTDB R214)*** | | | ***Known species fraction (Supplemented)*** | | | ***Cultured species fraction (GTDB R214)*** | | | ***Number of samples*** |
|---|---|---|---|---|---|---|---|---|---|---|---|
| ***Class*** | ***Biome*** | ***Mean (%)*** | | ***Median (%)*** | ***Mean (%)*** | | ***Median (%)*** | ***Mean (%)*** | | ***Median (%)*** | |
| *Eukaryotic host-associated* | Human | 81.05 | ± 17.95 | 86.34 | 80.02 | ± 17.26 | 84.80 | 51.22 | ± 24.69 | 52.46 | 76, 353 |
| | Mouse | 84.00 | ± 10.58 | 85.80 | 84.77 | ± 9.94 | 86.49 | 25.55 | ± 19.23 | 18.75 | 5, 767 |
| | Pig | 69.88 | ± 10.56 | 69.50 | 70.57 | ± 10.51 | 70.46 | 22.04 | ± 14.64 | 20.19 | 3, 446 |
| | Bovine | 45.66 | ± 19.59 | 43.40 | 48.51 | ± 19.80 | 48.62 | 4.88 | ± 10.70 | 1.36 | 1, 266 |
| | Plant | 53.43 | ± 38.34 | 66.21 | 55.62 | ± 36.14 | 67.72 | 43.37 | ± 36.57 | 39.17 | 833 |
| | **Subtotal** | **78.27** | **± 20.95** | **85.03** | **77.83** | **± 19.78** | **83.85** | **46.92** | **± 26.89** | **48.12** | **115, 575** |

| | | | | | | | | | | | |
|---|---|---|---|---|---|---|---|---|---|---|---|
| *Environmental* | Marine | 37.02 | ± 20.94 | 35.33 | 41.09 | ± 21.90 | 39.68 | 9.58 | ± 14.02 | 4.70 | 9, 768 |
| | Soil | 8.98 | ± 14.61 | 3.96 | 14.05 | ± 16.98 | 8.49 | 3.51 | ± 9.58 | 0.77 | 7, 951 |
| | Sediment | 17.46 | ± 21.72 | 8.92 | 20.04 | ± 22.43 | 11.75 | 5.51 | ± 11.15 | 1.19 | 3, 296 |
| | Freshwater | 37.22 | ± 20.71 | 38.21 | 44.96 | ± 21.86 | 46.28 | 10.44 | ± 14.34 | 5.71 | 2, 484 |
| | Aquatic | 33.36 | ± 26.78 | 28.18 | 39.95 | ± 27.05 | 36.09 | 15.53 | ± 23.86 | 5.80 | 1, 590 |
| | **Subtotal** | **25.56** | **± 23.77** | **19.85** | **30.24** | **± 24.85** | **24.99** | **8.01** | **± 14.80** | **2.57** | **25, 541** |
| *Misc* | Wastewater | 44.31 | ± 20.45 | 44.60 | 48.81 | ± 19.04 | 49.44 | 22.36 | ± 18.25 | 19.71 | 2, 691 |
| **Total** | | **64.42** | **± 31.64** | **78.75** | **65.88** | **± 29.41** | **78.09** | **36.62** | **± 30.06** | **34.60** | **167, 116** |

**Supplementary Table 2.**

Proportion of SingleM profile abundances that were classified at each taxonomic resolution in a SingleM taxonomic profile generated using a 58/60 bp (corresponding to ~97% ANI) or a 57/60 bp (corresponding to 95% ANI) cutoff for species assignment. Profiles with incorrect taxonomy are labelled "incorrect" and were nearly all species-level discrepancies. Genomes were chosen from genera with at least 3 species that each had at least 3 genomes in GTDB R214.

| Type | Taxonomy resolution | Proportion of OTU sequences | |
|---|---|---|---|
| | | 58bp (~97% cutoff) | 57bp (95% cutoff) |
| Sensitivity (correct species is in database) | domain | 0.1% | 0.1% |
| | phylum | 0.0% | 0.0% |
| | class | 0.1% | 0.1% |
| | order | 0.1% | 0.2% |
| | family | 0.5% | 0.5% |
| | genus | 5.0% | 4.1% |
| | correct species | 93.1% | 93.8% |
| | incorrect species | 1.1% | 1.1% |
| Specificity (correct species is missing from database) | domain | 0.4% | 0.3% |
| | phylum | 0.0% | 0.1% |
| | class | 0.3% | 0.5% |
| | order | 0.3% | 0.8% |
| | family | 3.0% | 4.3% |
| | genus | 76.3% | 68.2% |
| | correct species | N/A | N/A |

| | | | |
|---|---|---|---|
| | incorrect species | 19.7% | 25.8% |
